# Characterisation of mesenchymal stromal cells in the skin of Atlantic salmon

**DOI:** 10.1101/2024.02.23.581759

**Authors:** R. Ruiz Daniels, S.J. Salisbury, L. Sveen, R.S Taylor, M. Vaadal, T. Tengs, S.J. Monaghan, P.R. Villamayor, M. Ballantyne, C. Penaloza, M.D. Fast, J.E. Bron, R. Houston, N. Robinson, D. Robledo

## Abstract

**Background:** The skin serves as the first line of defence for an organism against the external environment. Despite the global significance of salmon in aquaculture, a critical component of this first line of defence, mesenchymal stromal cells, remains unexplored. These pluripotent cells can differentiate into various tissues, including bone, cartilage, tendon, ligament, adipocytes, dermis, muscle and connective tissue within the skin. These cells are pivotal for preserving the integrity of skin tissue throughout an organism’s lifespan and actively participate in wound healing processes.

**Results:** In this study, we characterise mesenchymal stromal cells in detail for the first time in healthy Atlantic salmon tissue and during the wound healing process. Single-nucleus sequencing and spatial transcriptomics revealed the transcriptional dynamics of these cells, elucidating the differentiation pathways leading to osteogenic and fibroblast lineages in the skin of Atlantic salmon. We charted their activity during an in vivo wound healing time course, showing clear evidence of their active role during this process, as they become transcriptionally more active during the remodelling stage of wound healing.

**Conclusions:** For the first time, we chart the activity of sub-clusters of differentiating stromal cells during the process of wound healing, revealing different spatial niches of the various MSC subclusters, and setting the stage for investigations into the manipulation of MSCs to improve fish health.

## BACKGROUND

Atlantic salmon is one of the most important aquaculture species worldwide, accounting for 32.6% of the global cultured marine finfish trade in 2020 (1), and with demand steadily increasing. However, sustainable expansion has been hindered by a range of disease and welfare issues. In particular, skin health issues, such as injuries caused by delousing operations, sea louse infections, and bacterial winter wounds, are on the rise and represent one of the main current challenges in Atlantic salmon aquaculture (2). These issues not only have major impacts on economic performance but also raise concerns about animal welfare.

The skin is a vital tissue for overall animal health, welfare, and robustness. In Atlantic salmon (*Salmo salar*, Linnaeus 1758), the skin is a complex organ that serves various functions, including protection against pathogens, mechanical stress, and homeostasis regulation. A key characteristic of fish skin is its remarkable ability to heal, which is enabled by specialized cells working in concert to repair and restore tissue integrity. Stromal cells, which are essential for maintaining various tissue structures, play a pivotal role not only during healing but also in homeostasis.

Mesenchymal stromal cells (MSCs) are fibroblast-like morphology that are present in various mammalian tissues (3–5) and are characterized by the expression of specific markers (e.g., CD34 and ITGA) and their ability to differentiate into adipocytes, chondrocytes, osteoblasts, and myocytes (6,7). In fish, MSCs have been identified in the fins of zebrafish (8), and their differentiation into chondrocytes, osteoblasts, and MSC-like cells has also been described in the visceral fat of Atlantic salmon (9). A better understanding of the ability of these cells to differentiate into myriad cell types and their role in the maintenance of tissue physiology and homeostasis can both provide insights into the mechanisms underlying skin repair and pave the way for a range of therapies aimed at improving wound healing (10).

The advent of single-cell technologies, such as single-cell RNA sequencing (scRNA-seq), has revolutionized our ability to study individual cells. While the extraction of individual cells can be challenging in certain tissues, such as the skin and fins (11), leading to biases in cell-type composition, this can be addressed through single-nuclei RNA sequencing (snRNA-seq), allowing the dissociation of nuclei from frozen tissue without losing cellular diversity (12,13). While single-cell/nuclear technologies reveal the transcriptome of individual cells, the spatial location of the cells is lost during dissociation. Combining snRNA-seq with technologies that capture the positional context of gene expression, such as spatial transcriptomics (14), allows the contextualisation of single-cell transcriptomic data. The power of combining these technologies to understand skin biology has already been exploited in humans, for example, to unravel the role of different skin cell types during wound healing processes (15). Although both single-nucleus RNA-seq and spatial transcriptomics have been used separately to study skin in Atlantic salmon (16,17), cell expression patterns have not yet been investigated in their tissue context, which is crucial to gaining understanding of their function.

This study aimed to advance our understanding of MSCs in Atlantic salmon. Using snRNA-seq of nuclei isolated from body skin (containing scales) and fins (lacking scales), we aimed to i) identify differences in cellular composition between body skin and fins, ii) identify mesenchymal stromal cells in Atlantic salmon skin, iii) infer their heterogeneity and trajectory dynamics and provide marker genes for these cells, and finally, iv) employ spatial transcriptomics to infer their activity during wound healing.

## RESULTS

Six snRNA-Seq Atlantic salmon skin and fin libraries were sequenced (two skin samples, two pelvic fins, one pectoral fin, and one dorsal fin). After quality control, 27,989 nuclei remained, averaging 4,665 nuclei per sample, with an average of 2,813 (± 791) unique transcripts (unique molecular identifiers - UMIs) and 1,566 (± 334) genes per nucleus. Summary statistics for all six libraries analysed can be found in Supplementary Table 1.

Unsupervised clustering of these 27,989 nuclei based on their transcriptomes revealed 18 distinct cell clusters (Figure 1A), potentially representing 18 different cell types. To assign putative cell types to the identified clusters, we referred to lists of marker genes from previously published works on the same tissues (16).

**Figure 1.**
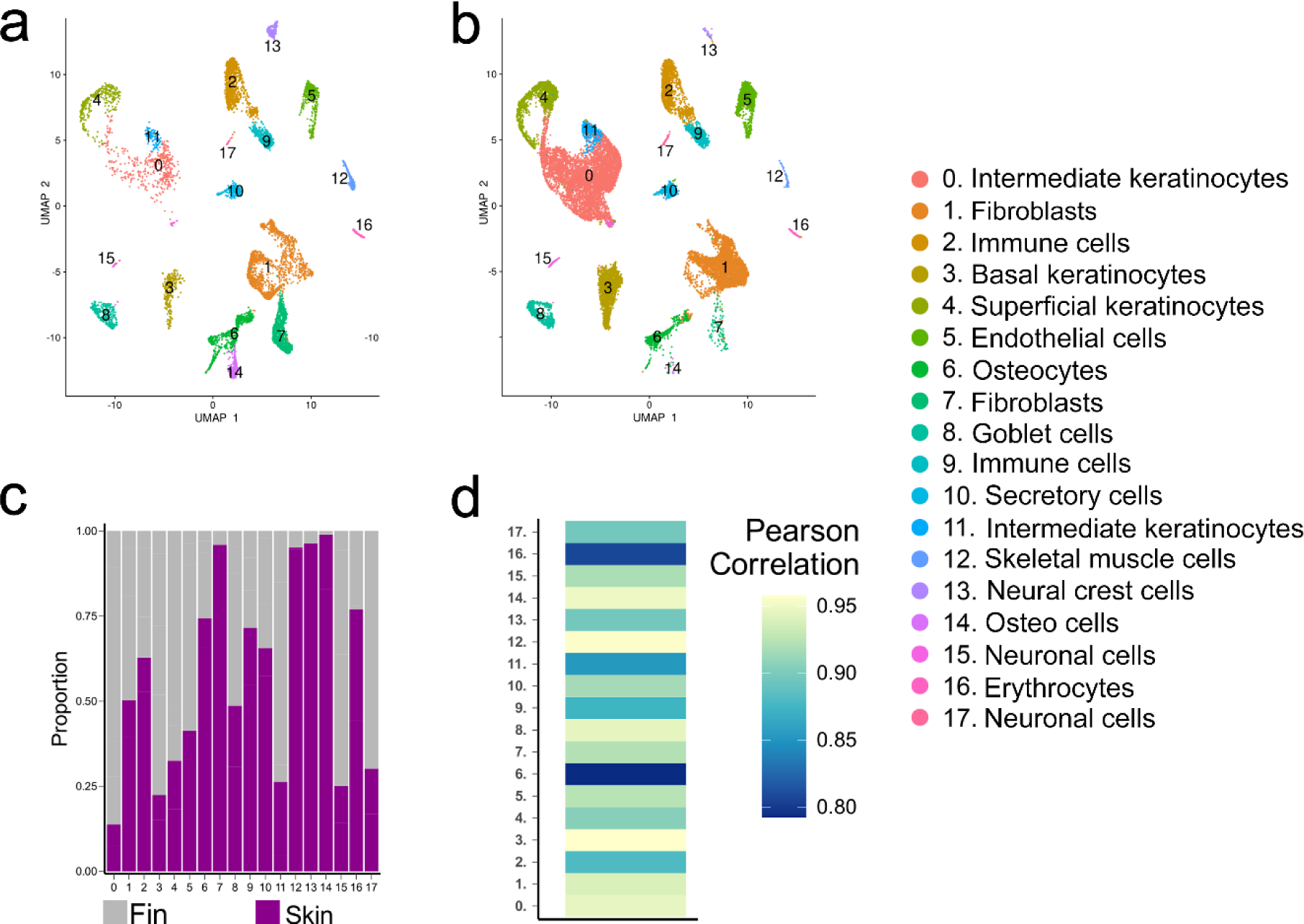
Comparison of cell types between fins and skin. A) Uniform manifold approximation and projection (UMAP) plot showing the 18 cell types present across all skin and B) fin samples. Cells are coloured according to their cluster, and the putative cell types are provided in the legend. C) Proportion of each cell type in the fin and skin normalized by dividing the number of cells of that cell type by the total number of cells. D) Pearson rank correlation between the transcriptomes of each cell type in fins and skin.

### Comparison of cellular composition between skin and fins

Although the skin covering the body (Fig. 2a) and the skin covering the fin (Fig. 2b) have broadly the same cell types, the proportions of some of these cell types differ between the fin and the body skin (Fig. 2c). Body skin showed a greater proportion of one population of fibroblasts (cluster 7), skeletal muscle cells (cluster 12), and neural crest cells (cluster 13), while the fin showed a greater proportion of intermediate keratinocytes (cluster 0), basal keratinocytes (cluster 3), another two populations of fibroblasts (clusters 1 and 11), and neural cells (cluster 17) (Figure 2c).

**Figure 2.**
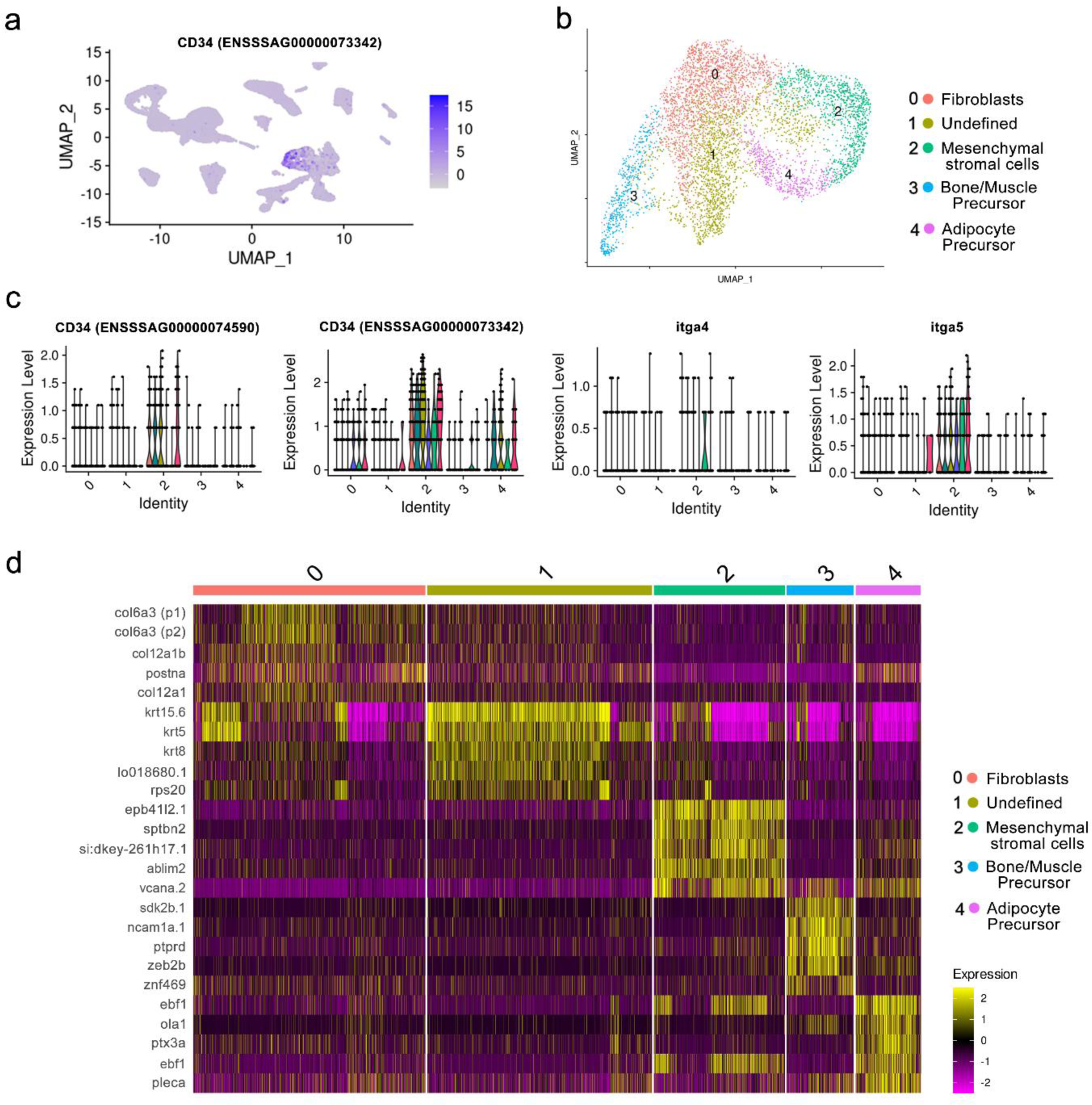
Identification and characterisation of mesenchymal stromal cells. A) UMAP showing the expression of known MSC markers such as *cd34* in salmon skin and fin cell atlas; expression of *cd34* (one paralogue). B) UMAP analysis of the MSC-containing population of fibroblasts. C) Violin plot showing the expression of MSC markers across the different subclusters of the fibroblast population. D) Heatmap showing the expression of the top 5 marker genes for each cluster for each tissue sample from left to right: fin1, fin2, skin1, fin3, fin4 and skin2 (supplementary Table 6 for the marker list).

### Fish mesenchymal stromal cells

To determine whether mesenchymal stromal cells are present in the skin and fins of Atlantic salmon fish, we used markers of this cell type identified in other species, specifically *cd34* (18) (Figure 2A), integrin α4 (*ITGA4*) (19), and integrin α5 (*ITGA5*) (20) (Supplementary Figure 3A). These genes were found to be expressed in a fibroblast population (cluster 1, Figure 3A). This cell population was reanalysed separately to explore potential heterogeneity within this cell type. Five subclusters were identified within this population (Figure 3B). Subcluster 2 included mesenchymal stromal cells with high expression of *itga4*, *itga5* and *cd34* (Figure 3C). The remaining clusters were identified based on their marker genes (Figure 3D) as fibroblasts (subcluster 0; expression of collagen and keratin genes), unclassified cells (subcluster 1; expression of nonspecific keratin genes), and two groups of differentiating MSCs (subclusters 3 and 4; expression of *fgfr4* and *wnt7*). Based on the expression of *pth1r*, *ptprd*, *runx3*, *fgfr4* and 3 bone/muscle precursors and based on the expression of *wnt7bb*, *ebf1*, *tafa5,* and *myh10,* we classified 4 as adipocyte precursors. See Supplementary Table 6 for a full list of markers and Table 1 for further classification and reference.

**Figure 3.**
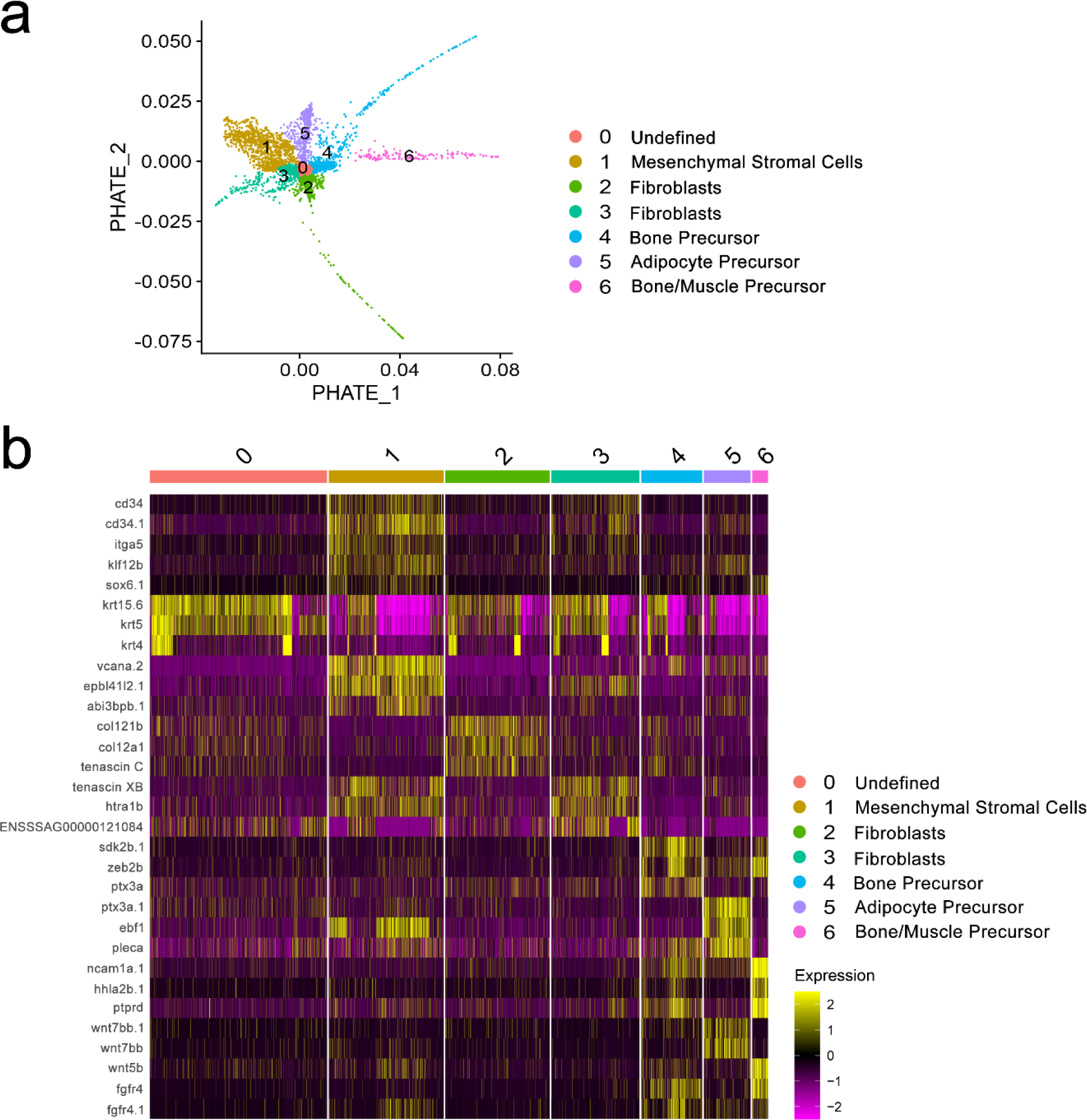
Identification and characterisation of pluripotent mesenchymal stromal cells using PHATE embedding. A) PHATE analysis showing the various states of the MSC-containing fibroblast cluster. B) Heatmap showing the top marker genes for the seven identified subcell states in MSC- containing fibroblast cluster 1. More details of these markers are provided in Table 1.

**Table 1.**
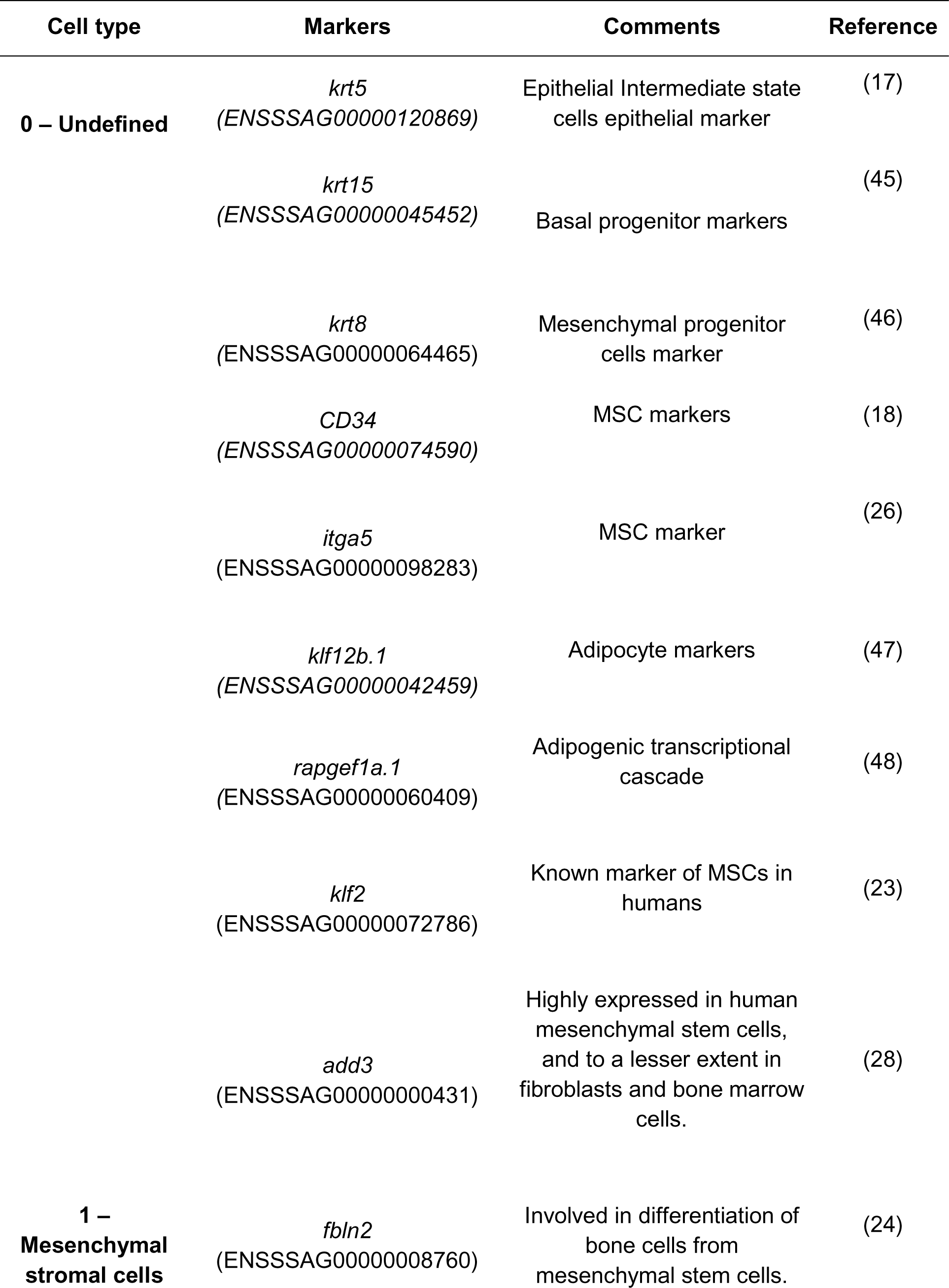
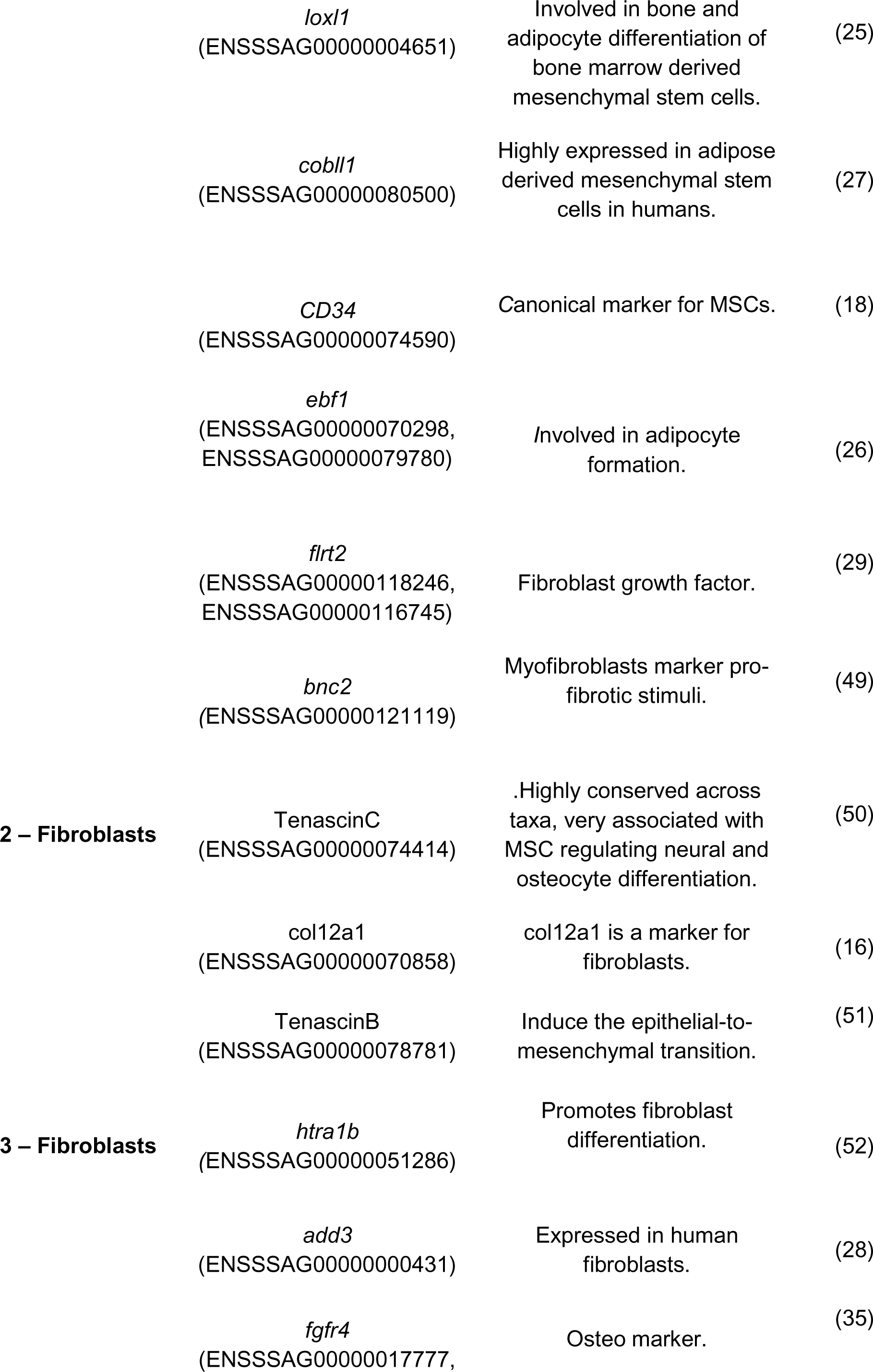
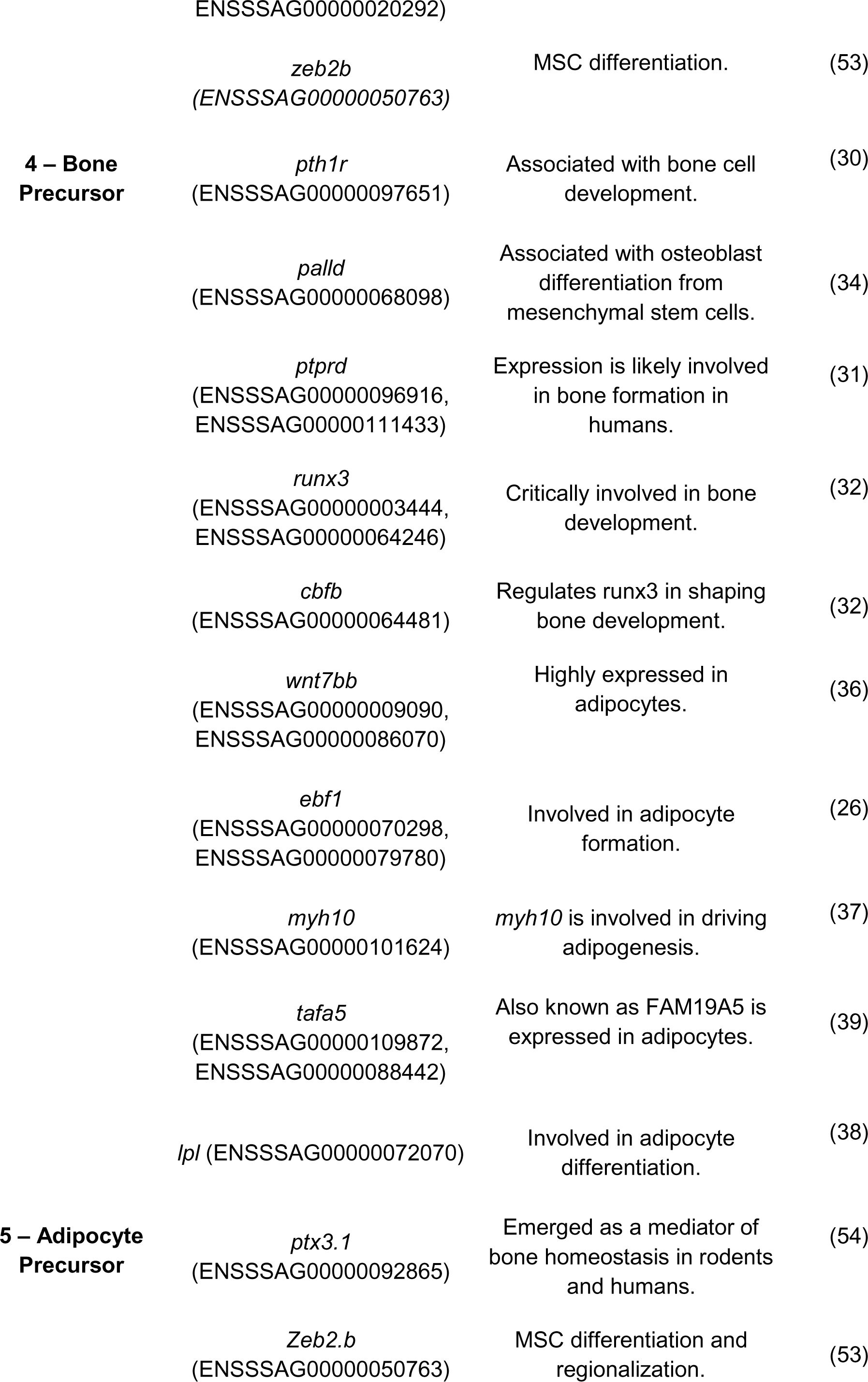
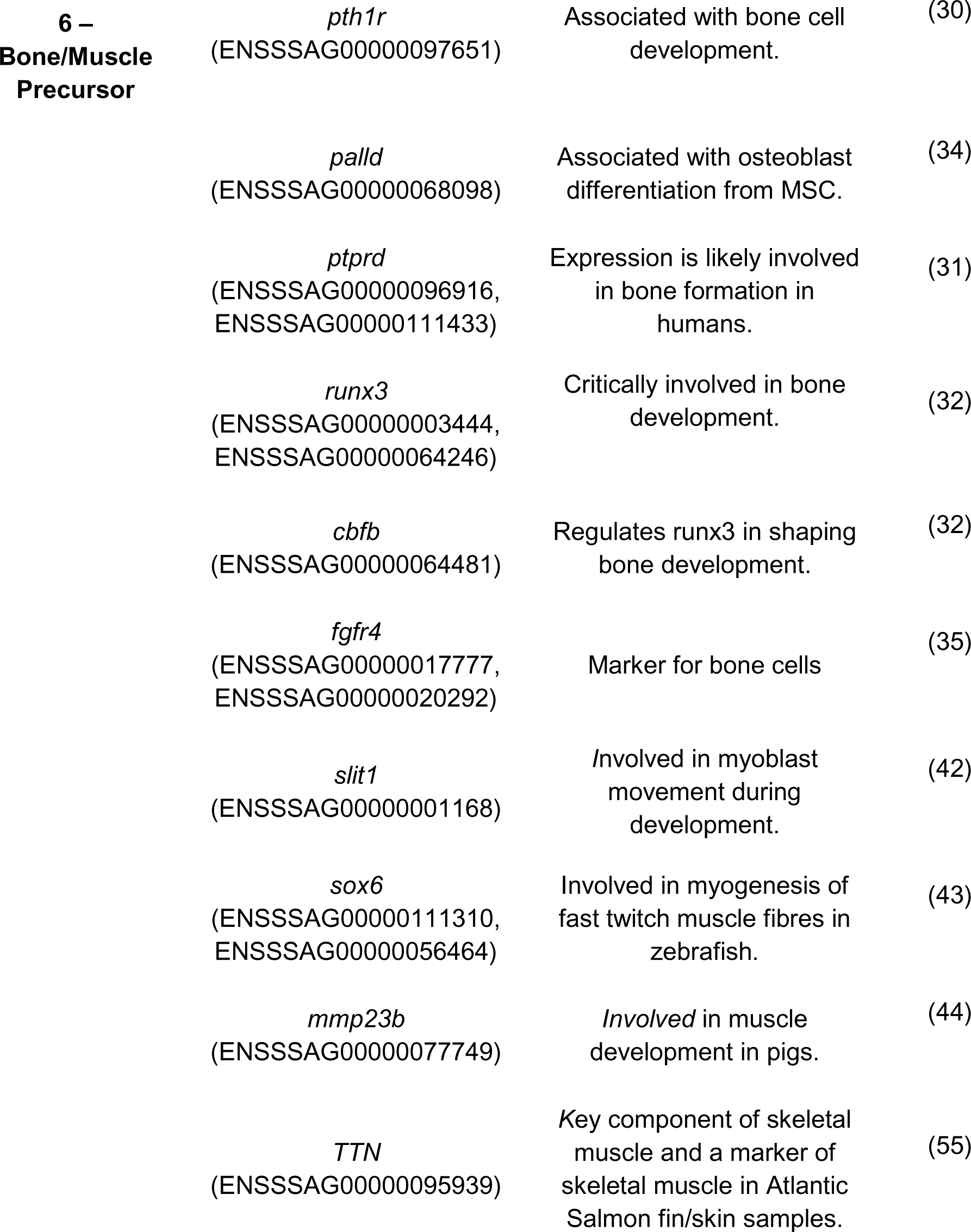
Cell states identified by PHATE in the MSC-containing fibroblast population and their marker genes.

### The dynamics of differentiating salmon mesenchymal stromal cells

UMAPs are useful visualization tools but are known to have limitations in faithfully capturing the dynamics and structure of high-dimensional data (21). To further explore the transitions of cell types, we used PHATE as an alternative dimensionality reduction method in the analysis of fibroblast cluster 1. PHATE identified 6 different cell subclusters (Figure 3a) with differential expression of marker genes (Figure 3b), 2 more than that observed in UMAP. This extra clustering reflects the transition probabilities between cell states, thus capturing more variability in the data in low-dimensional space while removing noise and retaining both the global and local structure (22). The correspondence between the UMAP and PHATE clusters is shown in Supplementary Figure 3B, with population 1 representing the mesenchymal stromal cells (Figure 3b, Table 1). PHATE also revealed different transitioning states, represented by the spindly arms observed (populations 2, 4, 6 and, to a lesser extent, 3), with the length of the arm indicating the extent of differentiation/change in a particular cluster/subcluster of cells (Figure 3a). These transitioning states were characterized based on their gene markers (Figure 3b, Table 1) and were identified as fibroblasts (2, 3), bone cell precursor cells (4), adipose precursor cells (5), muscle/bone precursor cells (6), and an undefined cell cluster (0) given its lack of biologically informative markers. This cluster was therefore characterized as “undefined”. (Figure 3b, see Table 1 for a list of key marker genes for each cluster).

Phate cluster 1 included MSCs based on their expression of the canonical MSC markers *CD34* (18) and *itga5* (20). This cluster also expressed *klf2*, a known marker of human mesenchymal stem cells (23). Consistent with their role as progenitors or bone cells and adipocytes (9), this cluster was characterized by multiple marker genes associated with osteogenesis and adipogenesis. For example, *fbln2* (24) is expressed by MSCs differentiating into bone cells, as is *loxl1* (25), which is involved in bone and adipocyte differentiation of MSCs. This cluster, along with cluster 4 (putative adipocyte precursor cells), also highly expressed *ebf1*, which is involved in adipocyte differentiation (26). An additional marker gene for cluster 1 was *cobll1* (27), which is highly expressed in adipose-derived MSCs. The proportion of cells in cluster 1 (MSCs) was greater than that in cluster 3 (fibroblasts). This finding is consistent with previous observations that this gene is highly expressed in MSCs and to a lesser extent in fibroblasts (28). PHATE cluster 2 was identified as fibroblasts by the expression of *col12a1*, which was previously found to be a marker of fibroblasts in several salmonid species (16), as well as *flrt2* (29), a fibroblast growth factor. The identity of cluster 4 as a bone precursor cell type is suggested by its expression of many genes associated with bone development, including *pth1r* (30), *ptprd* (31), *runx3* (32,33), and *cbfb* (32). Other marker genes supporting this identity include *palld*, a gene associated with osteoblast differentiation from MSCs(34), and *fgfr4,* a marker for bone cells (35), particularly for osteoblasts in Atlantic salmon skin and fin samples (16). The identity of PHATE cluster 5 as an adipocyte precursor cell is suggested by its expression of multiple genes associated with adipogenesis, including *wnt7bb*,(36) (Figure 4C), *ebf1* (26), *myh10* (37), and *lpl* (38). This cluster also expresses *tafa5a*, which is highly expressed in adipocytes (39). This cluster is unlikely to contain fully differentiated adipocytes, however, given the expression of *TTN*, a key marker for muscle cells in salmonid skin samples (16). Intriguingly, *sulf1,* which has been observed to be downregulated during adipogenesis (40), likely due to its role in driving osteogenesis (41), was also a marker for this cluster. This potentially reflects that this cluster is an early adipocyte precursor. Cluster 6 expressed markers associated with both bone and muscle differentiation. This cluster shared many marker genes with cluster 4 (bone precursor cells), including *palld*, *fgfr4*, *cbfb*, *runx3*, *ptprd*, *pth1r* and wnt5b (35). However, this cluster was also uniquely defined by multiple marker genes associated with muscle development: *slit1* (42), *sox6* (43), *mmp23b* (44), and *TTN* (16). Given that cluster 6 was not apparent in skin samples (Fig. 4b) and that skeletal muscle cells were predominantly detected in skin but not in fin samples using snRNAseq (16), this cluster may represent a muscle precursor cell that is not apparent in fin samples due to the decreased abundance of skeletal muscle in this tissue. For more detailed marker information and ENSBL gene IDs, see Table 1.

**Figure 4.**
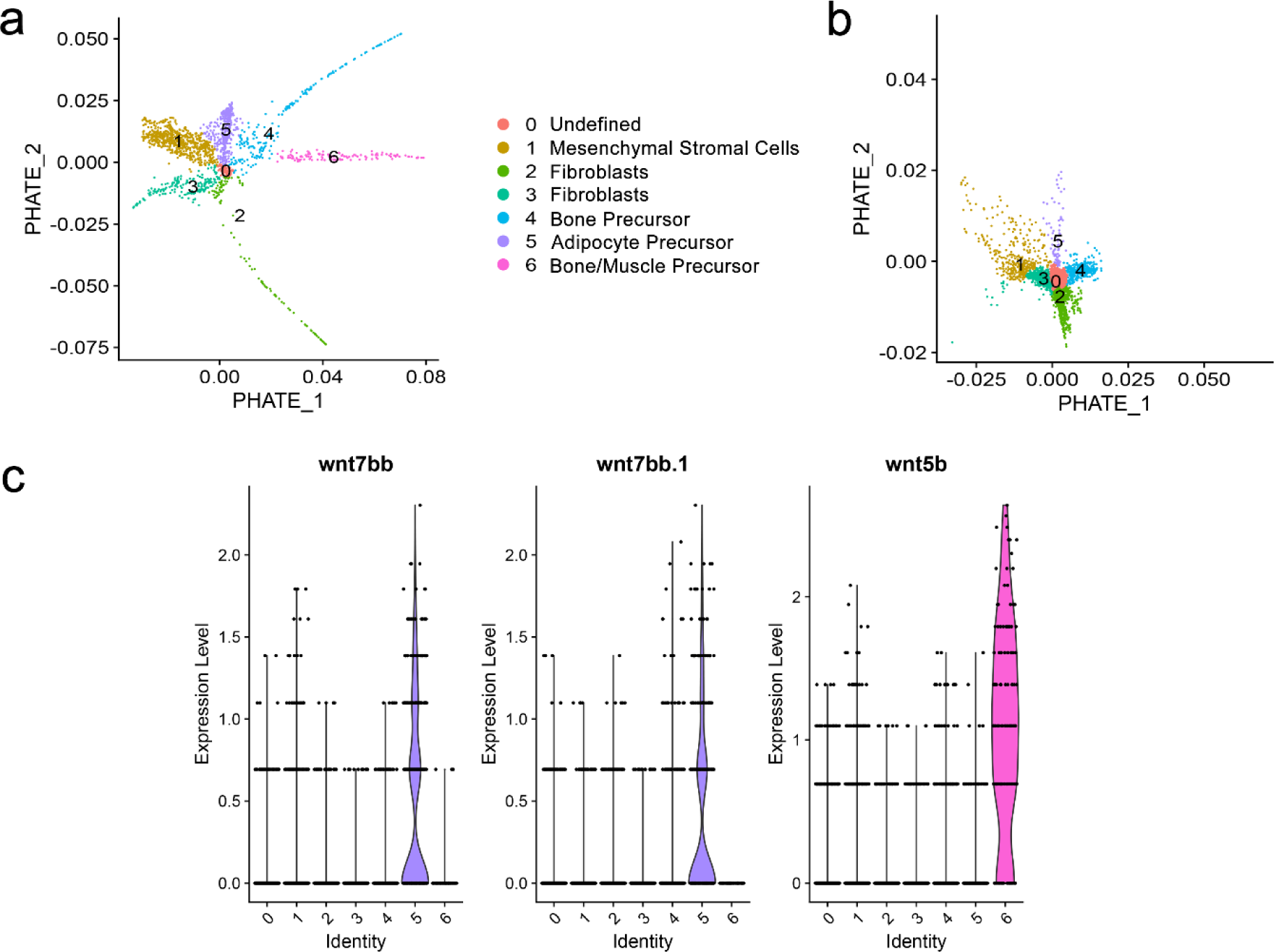
Differences in mesenchymal stromal cells between the skin and fin. A) PHATE plot showing the skin cells. B) PHATE plot showing the fin cells. C) Violin plot showing the expression of the *Wnt* genes in the 6 PHATE clusters.

### Differences in mesenchymal stromal cells in the fin and skin

The separate sub clustering of the MSC-containing fibroblast population for skin and fins showed clear differences between the two tissues (supplementary image 4 and supplementary figure 5, respectively). The skin showed a more complex structure than the fin, which is biologically consistent, as the skin has to regenerate a larger set of cell types, for example, scales and a fatty layer that is not present in fins. The differences became clearer when skin and fin cells were plotted separately (Figure 4a and b), with the fin presenting more quiescent cell subpopulations or states, especially in terms of (4) and bone/muscle (6) (Figure 4a & 4b). Fibroblast trajectories are also more pronounced in the skin. In general, this probably means that MSCs actively differentiate in the skin and are likely involved in general tissue homeostasis and regeneration, while the MSC population in the fins shows less proliferation and differentiation, but the population nevertheless still exists and is likely able to respond to external injuries and other factors affecting fin integrity.

### Mesenchymal stromal cells during wound healing

To investigate the activity of MSCs during wound healing, we used tissue sections of mechanically induced wounds at 2 and 14 days post wounding (dpw) (Figure 5a and 5b, respectively). These time points represent an early phase of wound healing (2 dpw), characterised by the formation of new epidermis, inflammation, and tissue degeneration (Figure 5a), and a late phase of wound healing (14 dpw) characterised by the replacement of damaged fibres with granulation tissue, a mix of new connective tissue and small blood vessels (Figure 5b). Using spatial transcriptomics, the expression of marker genes for MSCs (listed in Supplementary Table 7) during wound healing was assessed (proxies for MSC activity and proliferation). These markers exhibited a diffuse distribution at 2 dpw and were present throughout the whole tissue section with low expression, while at 14 dpw, expression/MSC activity was distinctly concentrated in the centre of the granulation tissue, displaying high levels of expression.

**Figure 5.**
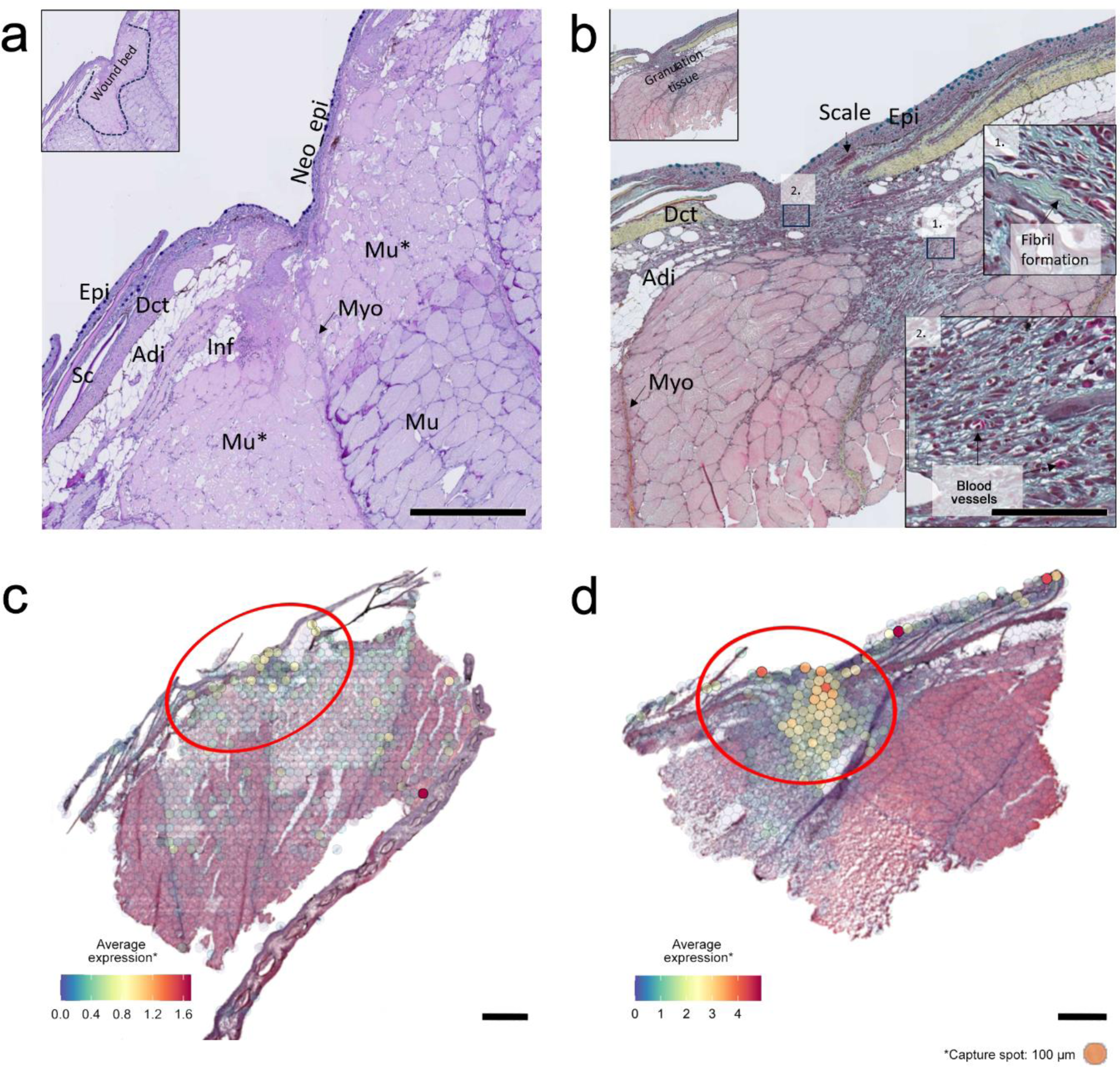
Wound healing in Atlantic salmon. Histological micrographs depicting incisional wounds at A) 2 days post wounding (dpw) (Alcian blue and PAS staining), the inflammation stage, and B) 14 dpw (Movat staining), the remodelling stage of wound healing. Epidermis (Epi), dense connective tissue (Dct), skeletal muscle fibres (Mu), damaged white muscle fibres (Mu*), myosepta (Myo), and newly formed epithelial tissue “Neo Epi”. Scales (Sc), adipose tissue (Adi), and polymorphonucleated inflammatory cells (InF). Expression of MSC-specific transcripts at C) 2 dpw and D) 14 dpw. Scale bars: 500 µm (a, b, c and d). (10X Visium spatial transcriptomic slides).

### Mesenchymal stromal cell subtypes during wound healing

To better explore the dynamics of MSCs at a finer scale during wound healing, the MSC subgroups identified during the PHATE embedding analysis (Figure 3) were plotted individually. To account for the transient state of the cells, the average expression of the top 20 most highly expressed genes was used (listed in Supplementary Table 7) for plotting on the spatial slides in Figure 6. Generally, all the MSC precursor subtypes exhibited lower average expression at 2 dpw (A, C, E, and G) than at 14 dpw (B, D, F, and H).

**Figure 6:**
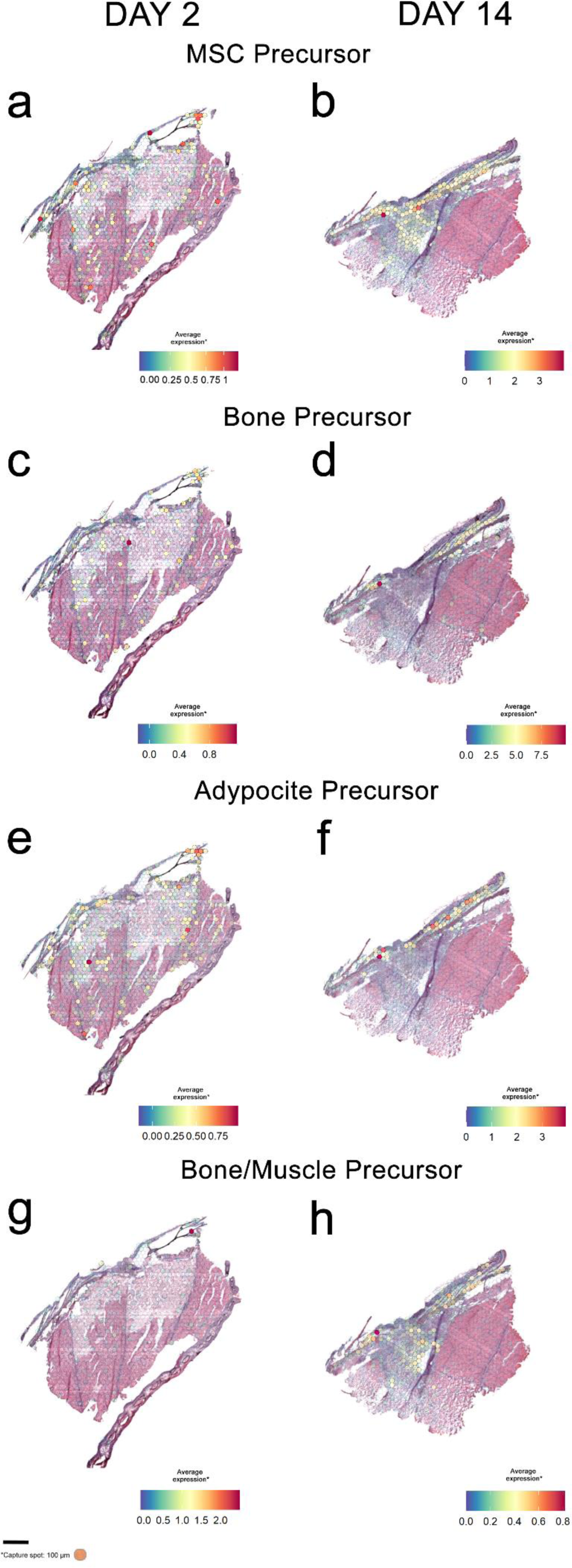
Charting the activity of different MSC subtypes. Expression of MSC subtype-specific transcripts, taken from the Phate analysis (average expression of the top 20 markers of each subtype, Supplementary Table 7), at 2 dpw (right) - the inflammation stage of wound healing, and 14 dpw (left) - the remodelling stage of wound healing. Scale bars: 500 µm (a, b, c, and d). (10X Visium Spatial Transcriptomic slides). A) Expression of MSC precursors on day 2 dpw. B) Expression of MSC precursors on day 14 dpw. C) Expression of bone precursors on day 2 dpw. D) Expression of bone precursors on day 14 dpw. E) Expression of adipocyte precursors on day 2 (dpw). F) Expression of adipocyte precursors on Day 14 dpw. G) Expression of bone/muscle precursors on Day 2 dpw. H) Expression of bone/muscle precursors on day 14 dpr.

The distribution of MSC precursors at 2 dpw was random across the section (A), but at 14 dpw, their distribution was more concentrated at the wound edge, granulation tissue, and layer of dense connective tissue (B). Similarly, with respect to the bone precursor, the same trend was observed, with low average expression at 2 dpw (C) and an increase in average expression at 14 dpw (D). However, there is less activity in the granulation tissue but high expression in the dense connective tissue, and scale pockets of this cell type do not seem to be active in the actual granulation tissue. There was random expression of adipocyte precursors across the section at 2 dpw (E), with an increase in expression at 14 dpw (F) in dense connective tissue, scale pockets, the apoptogenic layer, and the wound bed. The bone/muscle precursor, aside from one dot in the scale pockets at 2 dpw (G), shows low expression compared to that at 14 dpw (H), where the expression is quite widespread in the epidermis, area of dense connective tissue, and granulation tissue.

## DISCUSSION

This study provides a detailed analysis of the composition of Atlantic salmon skin and fins, focused on characterising mesenchymal stromal cells (MSCs) in this species. We identified MSCs among a population of fibroblasts using markers discovered in other species and revealed the multifaceted role of this cell population, which contains putative stem cells capable of differentiating into different fibroblast, osteocyte, muscle and adipocyte cell types. The integration of single-nucleus RNA-seq data and spatial transcriptomics revealed that these cells are present throughout the skin and play a major role in wound healing, especially during the tissue remodelling phase.

### Variation in the cellular composition of fins and skin

Fish skin and fins are generally very similar tissues. One of the main differences between the tissues is that the cell types annotated as “skeletal muscle”, “progenitor pigment cells” and “fibroblasts” are practically only present in the skin, likely reflecting the different anatomical architectures of the skin. Erythrocytes also show a relatively large transcriptomic divergence between the skin and fins. Erythrocytes in unlike in mammals, are nucleated and are also involved in immune responses (56), and perhaps the relatively low correlation between transcriptomes is simply a signature of greater transcriptional variation in this cell type.

However, the largest differences were observed in the osteo lineage, with osteocytes being more numerous in the skin and showing the greatest transcriptomic differences between the two tissues. MSCs in the skin seem to differentiate into two different osteo like cell types, putatively identified as bone precursors and bone muscle precursors, based on their gene expression and transition state. The bone precursors do not seem to be in such an active state in the fin compared to the scale, and the bone muscle precursors are not present in the fins. These differences could be due to bone regeneration in these tissues. The regeneration of scales in the skin is different from that of bones in the fin, as it is coordinated by the activity of osteoblasts (secretion and mineralization of the bone matrix) and osteoclasts (reabsorption of the bone matrix) to form an exoskeletal appendage. Bone cells also participate in the remodelling and vascularization of nerves and blood vessels within the scale tissue (57). The regeneration of bone in the fin is regulated by osteoblast migration and differentiation (58) into the area where the bone is regenerated. This work paves the way to further understanding how these cells are involved in the regeneration of scales and bones by providing markers to better understand the regeneration of bone in these two distinct bony structures.”

### Fish mesenchymal stromal cells

In this study, mesenchymal stromal cells (MSCs) were identified and characterised in Atlantic salmon. These MSCs are embedded in cell populations previously classified as fibroblasts (16) and show high expression of the two paralogues of *cd34*, *itga4* and *itga5*, which are known markers of MSCs in mammals (18–20). MSCs have been shown to be very similar to fibroblasts in morphology, proliferation dynamics, and immunomodulatory capacities, as well as in gene expression patterns (59), and some authors have even suggested that they are fibroblasts (60). Further exploration of this population of MSC-containing fibroblasts revealed signatures not only of stem cells but also of osteocytes. This suggested that MSCs could differentiate into various cell types and that this cell population was composed of stem cells, transitioning cell types and differentiated cell types. methods that take into account their transitioning nature to identify and characterize them. This is not unusual for MSCs, as their differentiation into various osteogenic and chondrogenic cell types has been described in humans (61). In Atlantic salmon, (9) described MSC-like cells isolated from fat tissue as highly plastic and undergoing transitions from osteogenic to adipogenic states. These findings indicate that MSCs in salmon are also likely to transition from a “pure” stromal cell state to various lineages, including bone and fat cells. This indicates that MSC-containing populations require more complex clustering.

The examination of the differentiation dynamics within this population revealed transitioning rather than static cell states, with clear cell differentiation niches in the form of “pure” MSCs and other transitioning MSC states, such as fibroblast, bone, adipocyte and bone muscle precursor cell states. Fish MSCs therefore appear to behave similarly to mammalian MSCs, putatively differentiating into other cell types when prompted. In theory, when they take a certain pathway, they should not be able to revert (e.g., fibroblasts vs. osteocytes) (4,9). Although our results cannot confirm whether these Atlantic salmon stem cells are transitioning from one state to another, the clear differentiation trajectories observed in our analyses suggest that this transition is unlikely.

### Wound healing

During the wound healing process, tissue damage necessitates replacement, and fish skin exhibits a remarkable regenerative ability. Zebrafish demonstrate complete scarless recovery from cutaneous wounds, even in adults (62). Fish, including salmon, employ mechanisms of wound closure similar to those in mammals, but they possess the unique capability of scarless regeneration throughout their lives, akin to embryonic mammals (63). Central to this phenomenon are MSCs, which play a crucial role in both wound healing and tissue regeneration (64). These versatile cells can differentiate into various essential cell types that are crucial for the regeneration of different tissues, such as connective tissue, bone, and cartilage. In this study, we detected MSC-specific markers at 2 days post wounding (dpw), but the distribution of these markers within the wound area and generally in the skin was scattered. In contrast, at 14 dpw, the markers for MSCs were notably concentrated within the wound bed. This concentration suggests potential mechanisms of cell proliferation and possibly even cell migration after the clearance of damaged fibres. The analysis revealed that MSCs are involved in the process of differentiation into various tissue types. For instance, bone and adipocytes and bone muscle share marker genes, as evidenced by the heatmap of the expression of the top 5 genes within the phate cluster (Figure 3c). An example is the gene *zeb2b*, which is involved in MSC differentiation and formation (53) and is expressed in clusters such as adipocytes 5 and bone 4. Other genes, such as different *Wnt* genes, are unique to each lineage (Figure 4c). This finding is consistent with previous reports showing that osteogenic and adipogenic cells share many common features, including transcriptomic signatures, but differentiate when they commit to one cell type or the other (9).

In this work, we explored the pivotal role that MSC subtypes play during the inflammation and remodelling phases of wound healing. MSCs are involved in cellular differentiation but also contribute to wound healing through various processes. For instance, they contribute to the secretion of essential proteins for the extracellular matrix of the skin, such as collagen, fibronectin or elastin (65). Moreover, MSCs exhibit immunomodulatory properties, influencing the immune response at the wound site by regulating immune cell activity. For instance, they contribute to the secretion of essential proteins for the extracellular matrix of the skin, such as collagen, fibronectin or elastin (66), thereby reducing excessive inflammation and promoting an environment conducive to tissue repair. To understand these different roles, MSC subtypes were explored during healing in vivo, and it was found that different subtypes of MSCs become active at different times during the healing phase. (Figure 6). For example, MSCs, bone, and adipocyte precursors are randomly expressed throughout the tissue section on day 2 of wound healing, with the bone muscle precursors being expressed at very low levels; however, their locations are very specific during the remodelling phase. The bone precursors are expressed in the scale beds around the wound granulation site, while the adipocyte precursors concentrate on the outer layer—the adipogenic layer—and become most highly expressed at the edge of the granulation site. The bone muscle precursors also showed this trend but with greater expression at the granulation site, similar to how MSC precursors concentrate at the granulation site, as previously observed (67). During day 14, the rebuilding of connective tissue started, requiring high MSC activity. There is also the beginning of scale formation, as indicated by the bone precursor cells, and adipogenesis starts at the granulation site. The bone muscle is also active. This study is the first to transcriptomically classify MSCs at the cellular level, characterize their differentiating niches, and demonstrate their various niches during the process of wound healing in a fish. This work paves the way for understanding stem cell differentiation at the transcriptomic level as well as at the spatiotemporal niche level.

### Potential applications of MSC knowledge in aquaculture

In humans, it has been shown that diet influences the differentiation and action of MSCs, leading to a variety of different health issues in multiple tissue types (68), and that MSC differentiation can shift toward adipogenic or osteogenic lineages by changing gene expression patterns (69). This opens an avenue for the exploration of nutritional modulation in salmon, which could have significant ramifications for fish health. Non healing skin wounds, including those caused by sea lice, are a constant issue in Atlantic salmon aquaculture, and new diseases such as winterulcer diseases caused by *Moritella viscosa* are becoming increasingly prevalent (2). Enhancing or manipulating MSC function may contribute to rapid and effective repair of the skin barrier, potentially mitigating the impact of skin damage and infections. These manipulations may also target other multifaceted contributions of MSCs, such as their role in promoting angiogenesis by stimulating the growth of capillaries, ensuring adequate blood supply, or their ability to secrete growth factors and cytokines, orchestrating the complex wound healing process, and facilitating tissue regeneration and recovery.

## CONCLUSION

This study advances our knowledge about the cellular composition of the skin and fins of Atlantic salmon and defines the activity of mesenchymal stromal cells in both tissues and during wound healing in a spatially resolved manner. Our study has described, for the first time, the fundamental dynamics of MSC differentiation in fish, with endpoints including fibroblasts, adipocytes, bone, and bone/muscle cells. The results also revealed the different spatial niches of the different MSC subtypes during the process of wound healing in fish skin. This finding sets the stage for future investigations into the potential manipulation of MSCs to improve fish health and welfare.

## METHODS

### Skin and fin sampling

The skin and fins of postsmolt Atlantic salmon (*Salmo salar* L.) were obtained from three different sources for single-cell nuclei extraction. Six Atlantic salmon with a mean weight of 486 g were randomly netted from a 2000 L stock tank at the Marine Environmental Research Laboratory (MERL) (Machrihanish, Scotland), University of Stirling. The fish were maintained at ambient sea temperature (14°C) in full-strength seawater (33%) in a flow-through system with a concentration of dissolved oxygen (DO) ranging between 8.6 and 8.8 ppm. The fish were fed commercial salmon pellets (Inicio Plus, BioMar, UK) at 1% of their body weight per day. The fish were culled by anaesthetic overdose using MS-222 (100 mg/L), and the brain was destroyed according to the methods of the UK Home Office Schedule 1. The fish lengths and weights were recorded immediately following euthanasia. Approximately 1 cm^3^ (1x1x1) samples of pectoral fin, dorsal fin, pelvic fin, and flank skin were dissected from the fish, immediately snap frozen in liquid nitrogen, and then stored at −70°C until further processing. Two additional Atlantic salmon with a mean weight of 25 ± X g were sampled from a recirculating system in the Center for Aquaculture Technologies (Prince Edward Island, Canada). The fish were placed in 135 L tanks with recirculating water at approximately 12°C. The fish were sedated using an anaesthetic dose of MS222 according to SOP/CATC/2085, followed by a lethal blow to the head. Fin and skin were collected in a cryotube, snap frozen on dry ice, and then stored at −70°C until further processing. One skin sample, one pelvic fin sample, and one pectoral fin sample from the first sampling and one skin sample, one pelvic fin sample, and one dorsal fin sample from the second sampling were selected for snRNA-seq.

### Nuclear isolation, library construction and sequencing

A tween with salt and tris buffer (TST) protocol adapted from (70) was used for nuclear extraction; this protocol had been previously optimized for Atlantic salmon fins and skin and is described in (71). Approximately 45 mg of flash-frozen skin or/fin was placed in a 6-well tissue culture plate (Stem Cell Technologies, cat. no. 38015) with 1 mL of TST buffer composed of 2 mL of 2X ST buffer, 120 µL of 1% Tween-20 (Sigma‒Aldrich, catalog no. P- 7949), 20 µL of 2% BSA (New England Biolabs, catalog no. B9000S), and 1.86 mL of nuclease-free water. The tissue was minced on ice for 10 minutes using Noyes Spring Scissors (Fine Science Tools, catalog no. 15514-12) for 10 minutes on ice. The resulting tissue homogenate was filtered through a 40 µm Falcon™ cell strainer (Thermo Fisher Scientific, catalog no. 08-771-2), and a further 1 mL of TST was used to wash the well and passed through the filter. The volume was increased to 5 mL with 3 mL of 1X ST buffer diluted from 2X ST buffer, which was composed of 146 µL of mM NaCl (Thermo Fisher Scientific, catalog no. AM9759), 292 µL of 10 mM Tris-HCl, pH 7.5 (Thermo Fisher Scientific, catalog no. 15567027), 10 µL of mM CaCl2 (Vwr, E506-100 mL), 210 µL of mM MgCl2 (Sigma–Aldrich, catalog no. M1028) and 9388 mL of nuclease-free water. The sample was then centrifuged at 4°C for 5 minutes at 500 g and 4°C in a swinging bucket centrifuge. The resulting pellet was resuspended in 1 mL of 958 PBS and 0.02% BSA buffer. The nucleus solution was then filtered through a 40 µm cell strainer (Falcon™). The TST and PBS buffers contained 200 U mL^-1^ of the ultrapure protector RNase inhibitor (Sigma‒Aldrich, catalog no. 3335399001), and the 1xST contained 20 U mL^-1^.

Salmon skin and fin nuclei were processed through the Chromium Single Cell Platform using the Chromium Single Cell 3’ Library and Gel Bead Kit v3.1 (10X Genomics, Pn-10001221) and the Chromium Single Cell A Chip Kit (10X Genomics, PN-120236) according to the manufacturer’s protocol. Briefly, after isolation, the nuclei were counted using a disposable hemocytometer (Neubauer C-Chip DHC-N01). The chromium was loaded with the aim of recovering 7,000 nuclei in a Chromium 3’Chip. The nuclei were then partitioned into droplets in the Chromium controller. Once the emulsions were formed, RNA was barcoded and reverse transcribed. The resulting cDNA was then amplified and fragmented, and adaptors and sample indices were added. Libraries were sequenced on a NovaSeq 6000 platform by Azenta, resulting in approximately 220 million paired-end 150 bp reads per sample.

### Genome Indexing and Read Alignment with STAR

Genome indexing and library mapping were performed with STAR (version 2.7.10a, (72). We appended the mitochondrial genome from the ENSEMBL V2 Atlantic salmon genome (Salmo_salar.ICSASG_v2.dna_rm.toplevel.fa.gz, v2, release 105, masked genome, assembly ID: GCA_000233375.4) to the ENSEMBL V3 Atlantic salmon genome (Salmo_salar.Ssal_v3.1.dna_rm.toplevel.fa.gz, v3.1, release 106, masked genome, assembly ID: GCA_905237065.2) for both the .gff and .fna files prior to indexing. The genome annotation .gff files were converted to .gtf files using gffread (v0.10.1) (73). The Atlantic salmon genome was indexed using STAR (--runMode genomeGenerate). Each library was then mapped against its genome with the 10X V3 cell barcode whitelist (3 M-February-2018.txt) using standard parameters for single-cell libraries (--soloMultiMappers Unique --soloBarcodeReadLength 150 --soloType CB_UMI_Simple --soloUMIlen 12 -- soloCBwhitelist 3 M-February-2018.txt --soloFeatures GeneFull --clipAdapterType CellRanger4 --outFilterScoreMin 30 --soloCBmatchWLtype 1MM_multi_Nbase_pseudocounts --soloUMIfiltering MultiGeneUMI_CR --soloUMIdedup 1MM_CR --readFilesCommand zcat --outSAMtype BAM Unsorted). The raw (unfiltered) files (genes.tsv, barcodes.tsv, and matrix.mtx) generated for each sample were then used for downstream analysis.

### Quality Control, Clustering, Integration

The raw STAR files were then analysed in the R (v4) environment using Seurat (v4.1, (74). Seurat objects for each library were created after removing nuclei with fewer than 200 features and features occurring in fewer than three nuclei. Nuclei where mtDNA features represented more than 10% of their total UMIs were removed, and then, mtDNA features were removed from the dataset (leaving 46832 features). Upper and lower thresholds for UMI and feature counts per nucleus were then determined for each sample based on knee plot visualization. For all samples, only nuclei with more than 500 UMIs but less than 6000 UMIs and more than 500 features and less than 3500 features were retained.

The samples were then merged into a single Seurat. The “v2” SCTransform version with the glmGamPoi method (v 1.8.0, Ahlmann-Eltze and Huber, 2021) was used to normalize RNA counts for each sample prior to calculating cell cycle scores using the “CellCycleScoring” function based on the “SCT” assay (see Supplementary Table 2 for list of genes used), regressing out scores for the S and G2M cell cycle stages. The SCTransformation was then repeated as described above, but additionally, the “S. Score” and “G2M. Score” variables.

Linear dimension reduction was conducted for each sample using the “RunPCA” function with 50 PCs. After consulting the elbow plots for each sample, a UMAP using 20 PCs was run for each sample, and the “FindNeighbours” function was applied using 20 PCs before using the “FindClusters” function with a resolution of 0.2. DoubletFinder (v 2.0.3, 76). was then applied independently to each sample by selecting the pK values with the highest associated BCmvn values. We assumed a 4% doublet formation rate (based on the Chromium instrument specifications) and adjusted for homotypic doublets (see Supplementary Table 3) for the remaining cells per sample after doublet removal.

The samples were integrated using 10,000 features, and the anchors were identified via the “rpca” reduction method and the “FindIntegrationAnchors” function. PCA was performed on the integrated dataset using 50 PCs, and 30 PCs were used for subsequent UMAP generation and clustering with a resolution of 0.2 (supplementary figure 1). Markers for each cluster were assessed using the logistic regression method and the FindAllMarkers function on the “SCT” assay and “data” slot, using sample ID as a latent variable to help reduce batch effects among samples. We used a pseudocount of 0.001, set a p value threshold of 0.01, and considered only genes that were upregulated, expressed in at least 25% of all nuclei (in either of the compared groups), and demonstrated the default threshold of 0.25 X difference (log-scale) between the two compared groups.

Four clusters were removed from the dataset due to the small number of distinguishing marker genes. Clusters 1, 3, and 7 also had very low average feature/UMI counts (Supplementary Figure 1 B). The SCTransformation was then repeated for each sample based on the RNA assay as described above, and the integration of samples for each species was conducted as described above using 30 PCs for UMAP generation and a resolution of 0.2 for clustering (see Supplementary Table 3 for the remaining cells per sample and Supplementary Figure 2 for the distribution of UMIs and features per sample and per cell type after all filtering).

### Comparison between the epidermis of skin and fins

To account for differences in sample sizes and sequencing, cell counts in each cluster were normalized by dividing the total number of cells of each cluster in that tissue type by the total number of cells captured in that tissue type. The transcriptomic similarity between tissues for each cell population was determined by Pearson rank correlation, using the mean expression of the set of all marker genes (Supplementary Table 4 contains the main representative markers, while Supplementary Table 5 includes the top 20 markers). Furthermore, (16) for a study of these clusters).

### Inference and dynamics of mesenchymal stromal cells

Known stem cell markers, specifically cd34 (18), integrin α5 (ITGA5) (20) and integrin alpha 6 (ITGA6) (77,78), were used to infer putative clusters that would contain mesenchymal stromal cells (MSCs). These populations were subset into a new Seurat object, normalized with SCTransform to generate updated residuals, and reclustered with the number of principal components informing the clustering and resolution chosen to provide biologically meaningful clusters (20 PCs and 0.3 resolution). PHATE (22) was used to generate two-dimensional embeddings to identify differentiation processes between cell types. PHATE embeddings were used to recluster the populations using the FindNeighbors and FindClusters Seurat commands.

### Wound healing samples and spatial transcriptomics

An Atlantic salmon wounding experiment was carried out as described by (67) with minor modifications. Briefly, the fish were fully anaesthetized with MS-222 (Sigma‒Aldrich). Incisional skin ulcers were introduced on the flank of the fish with a sterile scalpel blade, which was cut through all skin layers, resulting in deep cutaneous wounds with damaged muscle fibres. Samples (N = 2 fish per sampling point and group) were taken on days 2 and 14 post wounding at the wound site. Tissue samples were frozen immediately in liquid nitrogen and stored at −80 °C. The fish were euthanized with an overdose of anaesthetic (MS-222) prior to tissue sampling. The fish were in good health prior to incisional wounding, and there were no mortalities. Spatial transcriptomic libraries were prepared as described previously (17). Summary statistics for spatial transcriptomics libraries can be found in Supplementary Table 1. The expression of specific MSC markers was inferred using the snRNA-seq atlas. This was achieved by plotting the top markers across the entire cell atlas of snRNA-seq data and retaining only those expressed exclusively by the MSC cell population, which was not present in any other cell cluster across the atlas (Supplementary Table 8).

To map the different MSC subtypes onto the spatial slide and considering the coexpression of these cell types among the MSC subtypes, the top 20 expressed genes for each cell subtype (including those in Table 1) were taken from the PHATE output (Supplementary Table 7). The average expression of these 20 marker genes for each MSC cell subtype was plotted on a spatial slide.

## ADDITIONAL INFORMATION

### Data availability statement

The datasets presented in this study will be found in NCBI public online repositorie, upon acceptance in peer-reviewed publication.

## Funding

This work was supported by FHF grant 901631 (“CrispResist”), FHF grant 901656 (ERN samspill), BBSRC grants BB/V009818/1 and BB/V009990/1 (“GenoLice”) and BBSRC Institute Strategic Grants to the Roslin Institute (BBS/E/20002172, BBS/E/D/30002275, BBS/E/D/10002070 and BBS/E/RL/230002A). DR also acknowledges funding from the Axencia Galega the Innovación (GAIN, Xunta de Galicia) as part of the Oportunius programme. SJS gratefully acknowledges funding from an NSERC PDF award.

### Author contributions

*The* MDF, SJM, LS, JEB, MDF, RDH, NR, and DR obtained funding. RRD, MDF, SM and DR conducted the experiments and collected tissue samples. RRD led the lab work for the snRNA-seq with assistance from SJS, PRV, MB and CP. The snRNA-seq RRD led the data analysis and interpretation with the assistance of SJS; PHATE analysis with the assistance of RST; and figure generation with the assistance of DR, PRV and LS. Spatial transcriptomics work led by LS with assistance from MV, TG and RRD. RRD led the writing in consultation with DR and with assistance from SJM, JEB, PRV, MDF, LS, RDH, and NR. All authors approved the final manuscript.

### Ethical statement

All animals were in good health prior to enrolment in the study, and no mortalities were recorded. The use of animals for experiments was in accordance with the guidelines of the EU legislation (2010/63/U) as well as the Norwegian legislation on animal experimentation and was approved by the Norwegian Animal Research Authority (28346).

## Acknowledgements

The authors wish to thank Professor Bente Ruyter for sharing samples with the project and the personnel at Sunndalsøra Research Station for maintaining the fish. Additionally, the authors express gratitude to Chessor Matthew and David Bassett from the Marine Environmental Research Laboratory (MERL) for providing samples for the project. Special thanks also go to Dr. Mark Braceland and Dr. Haitham Mohammed for their contributions to providing samples.

## SUPPLEMENTARY FILES

**Supplementary Figure 1.**
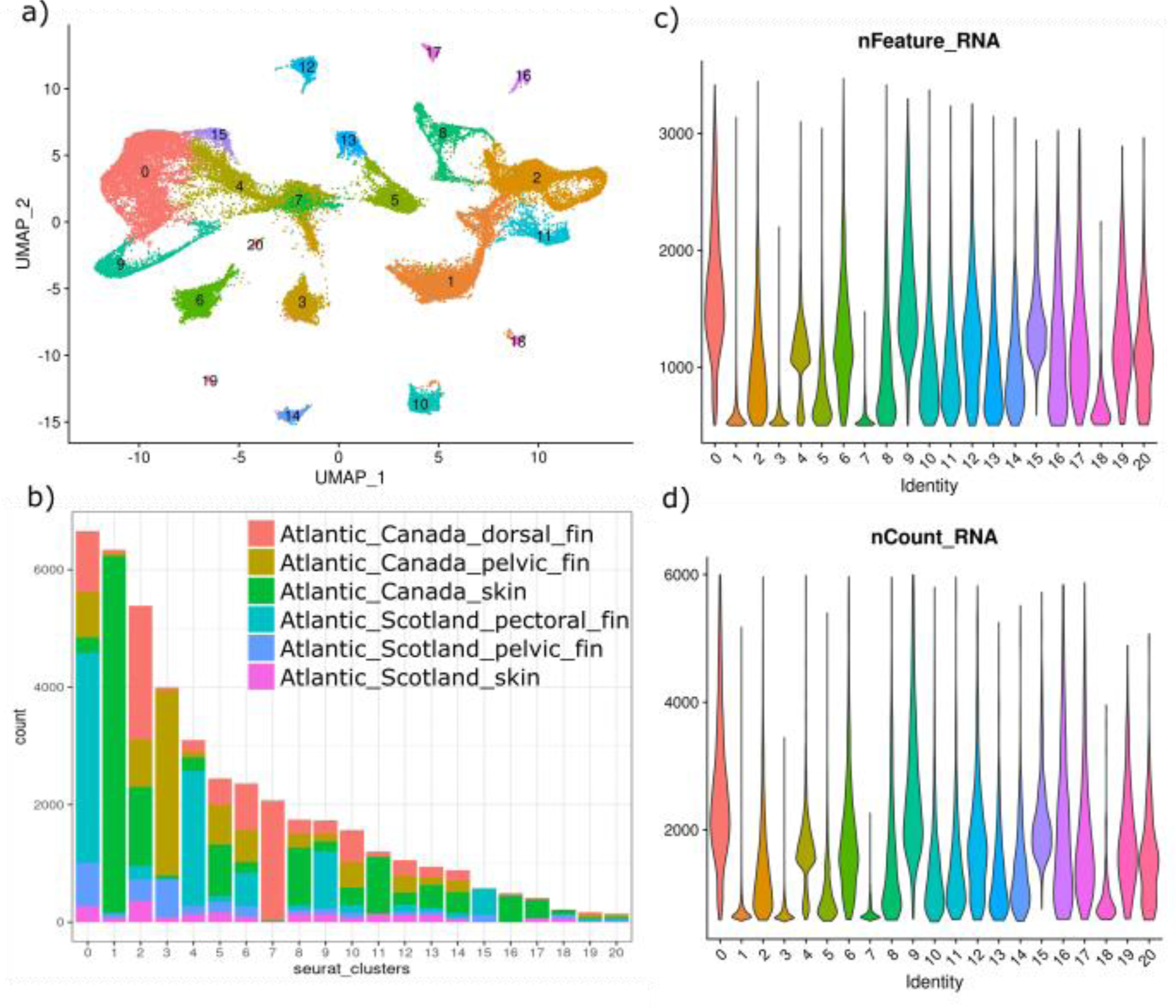
Cell clusters after initial integration of samples: a) UMAP, b) number of cells per cluster per sample, c) Violin plot of the distribution of feature counts per cluster, d) Violin plot of the distribution of UMI counts per cluster. Clusters 1, 3, 4, and 7 were removed from subsequent analysis. Clusters 1, 2, and 7 demonstrated low average UMI/feature counts. Clusters 1, 3, 4, and 7 were also found almost exclusively in a single sample.

**Supplementary figure 2.**
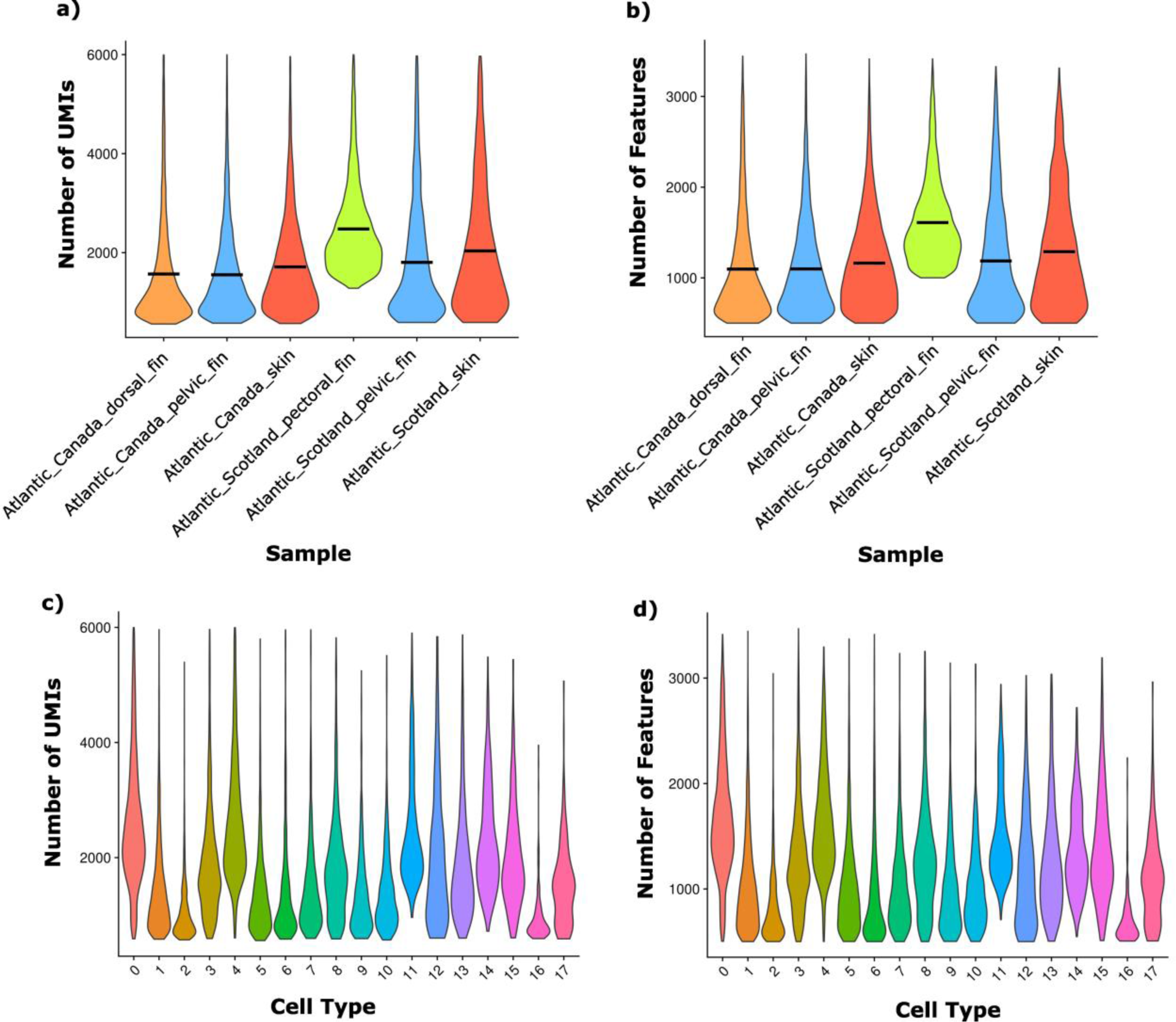
Distribution of UMIs (a) and features (b) for each sample after filtering (the horizontal black bar indicates the mean value for each sample). Distribution of UMIs (c) and features (d) for each cell type after filtering.

**Supplementary figure 3.**
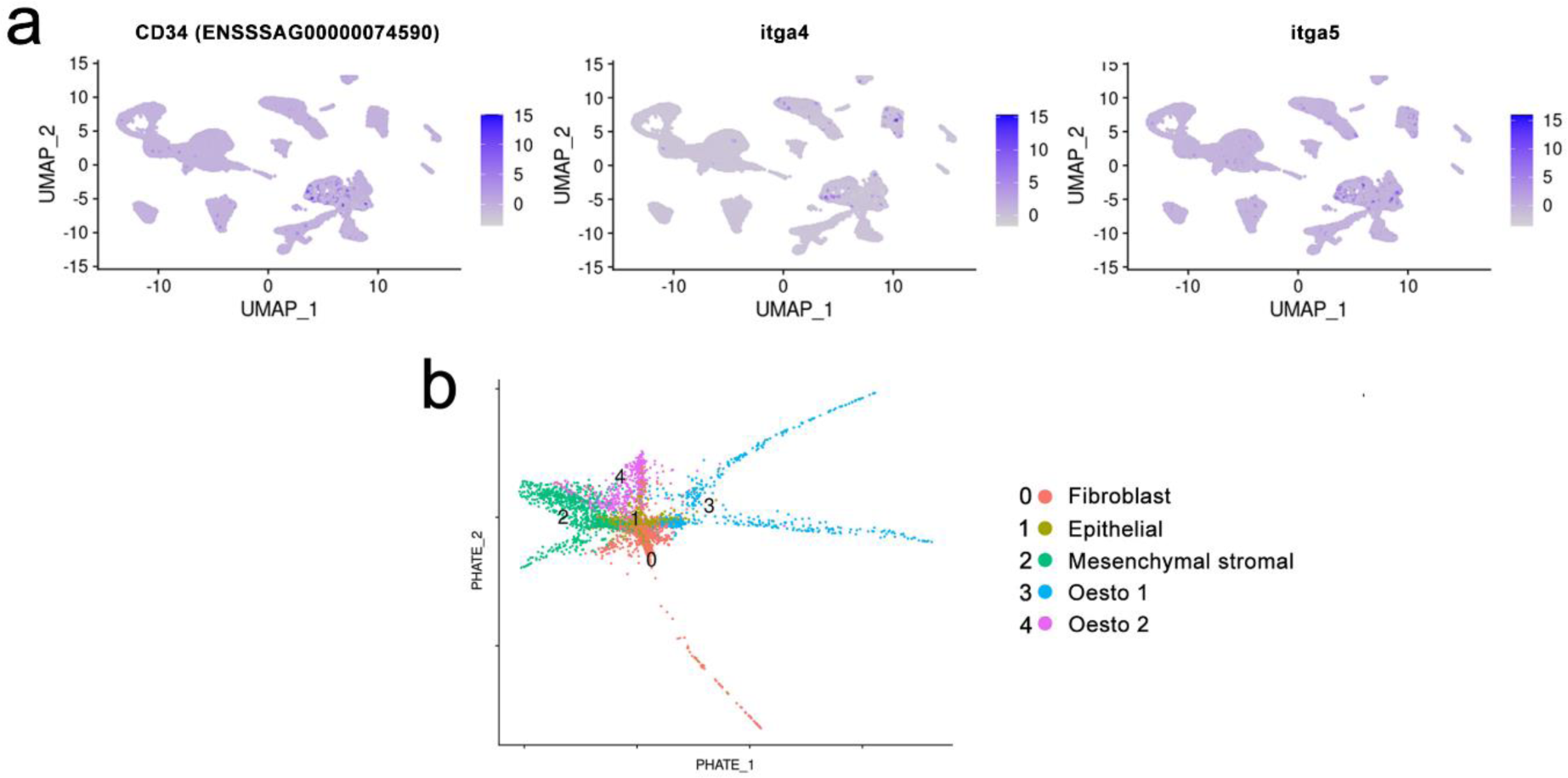
A) UMAP showing the expression of known MSC markers in salmon skin and in the fin cell atlas; the expression of the cd34 paralogue integrin α4 (itga4) and integrin alpha 5 (itga5) is shown. B) PHATE analysis of fibroblast cluster 1 (k=5) showing the differentiation dynamics of these cells; the clusters correspond with those of the Seurat analysis.

**Supplementary figure 4.**
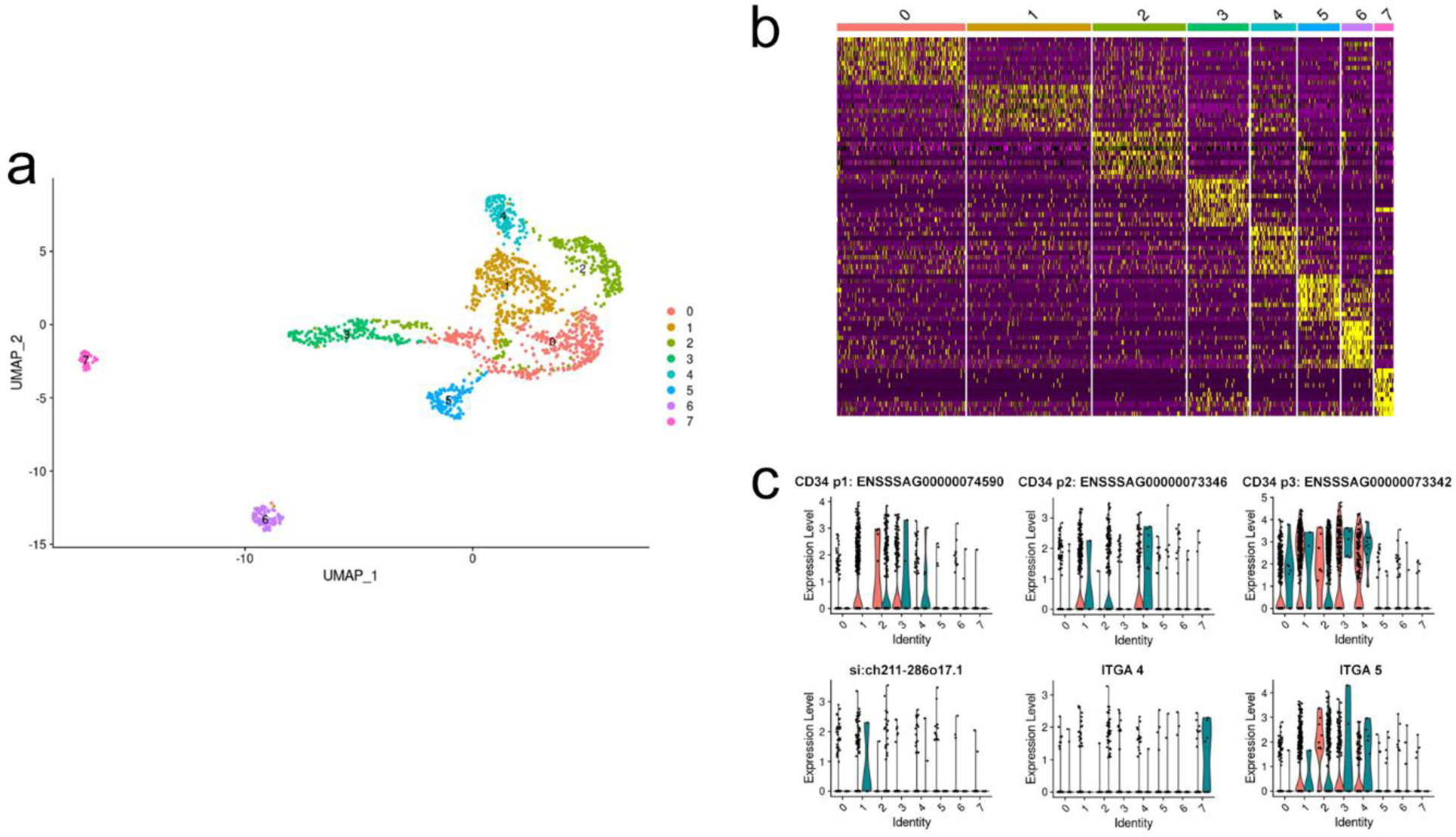
Mesenchymal stromal cells in the skin. a) Uniform manifold approximation and projection (UMAP) plot showing the 7 cell types present in all the scaly skin data for the fibroblasts that we could rename putative hemopoietic cells are colored by type and annotated to the right. Population 1. Fibroblasts (putative haematopoietic cells) were subsetted from the scaly skin data and renormalized using SC transformation, so the identities of the cell types are dependent only on the genes expressed within a cluster and are reclustered. A complete list of the markers is provided in Supplementary Table 9. b) Heatmap showing the gene expression signatures of each hematopoietic stem cell type. On the left, each column represents one cell type, and each row indicates the expression of one gene; the size of the dot represents the cell number, and the value for each gene is the row-scaled *Z* score. The top markers for each cell type are shown in Supplementary Table 9. c) Violin plot that combines the features of a box plot and a density plot. The distribution of stem-like transcripts across cell populations for each of the skin libraries is displayed. These findings indicate that populations 1, 2, 3 and 4 express stem cell markers.

**Supplementary figure 5.**
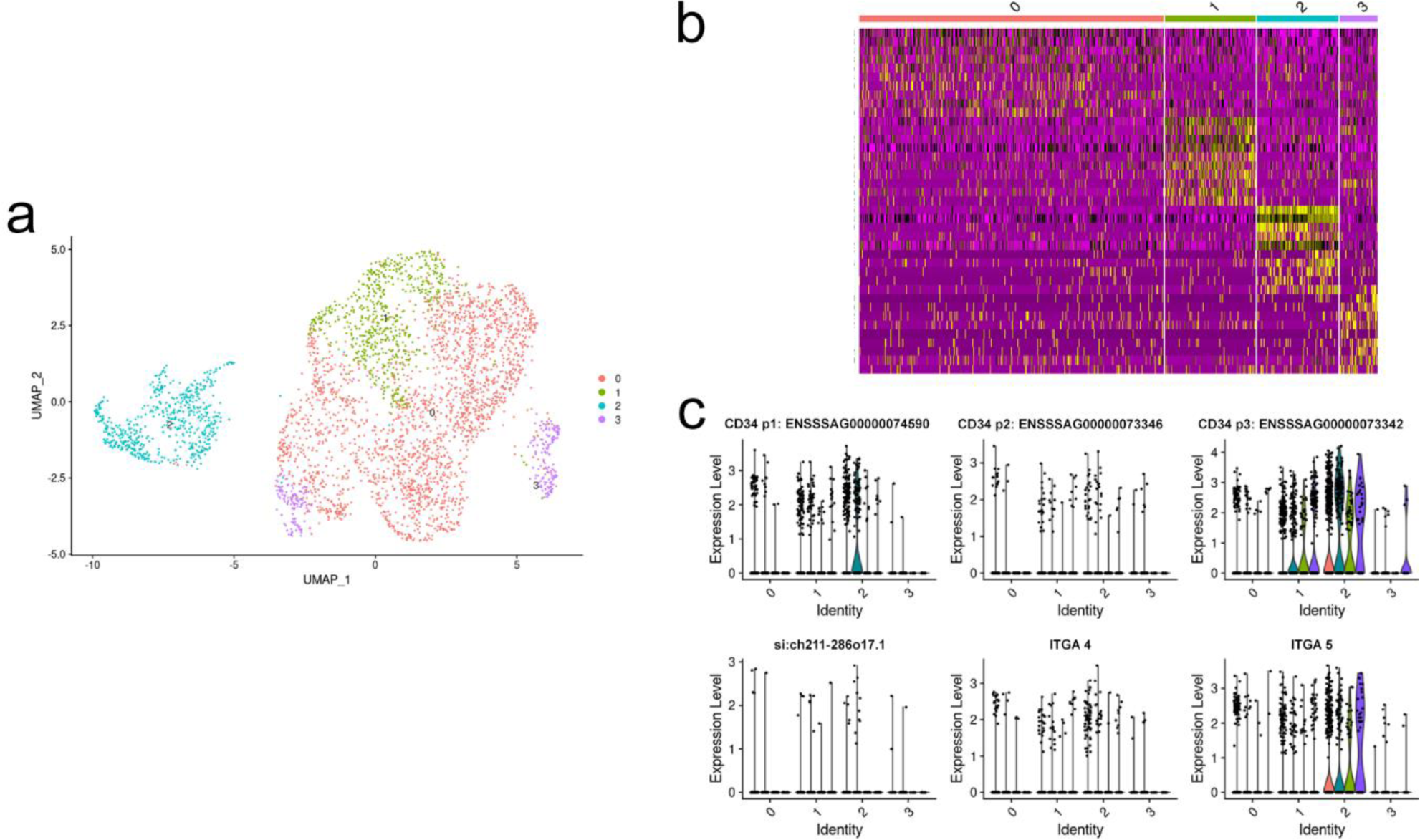
Mesenchymal stromal cells in the fin. a) Uniform manifold approximation and projection (UMAP) plot showing the 4 cell types present in all the scaly skin data for the mesenchymal stromal cells coloured by type and annotated to the right. The mesenchymal stem cell cluster was subsetted from the fin data and renormalized using SCtransform, so the identities of the cell types are dependent only on the genes expressed within a cluster and are reclustered. The complete list of markers is provided in Supplementary Table 10. d) Violin plot that combines the features of a box plot and a density plot. The distribution of stem-like transcripts across cell populations for each of the fin libraries is displayed. This indicates that populations 1 and 2 are putative stem cells. c) Heatmap showing the gene expression signatures of each hematopoietic stem cell type. On the left, each column represents one cell type, and each row indicates the expression of one gene; the size of the dot represents the cell number, and the value for each gene is the row- scaled Z score. The top markers for each cell type are shown in Supplementary Table 10.

**Supplementary Table 1.**
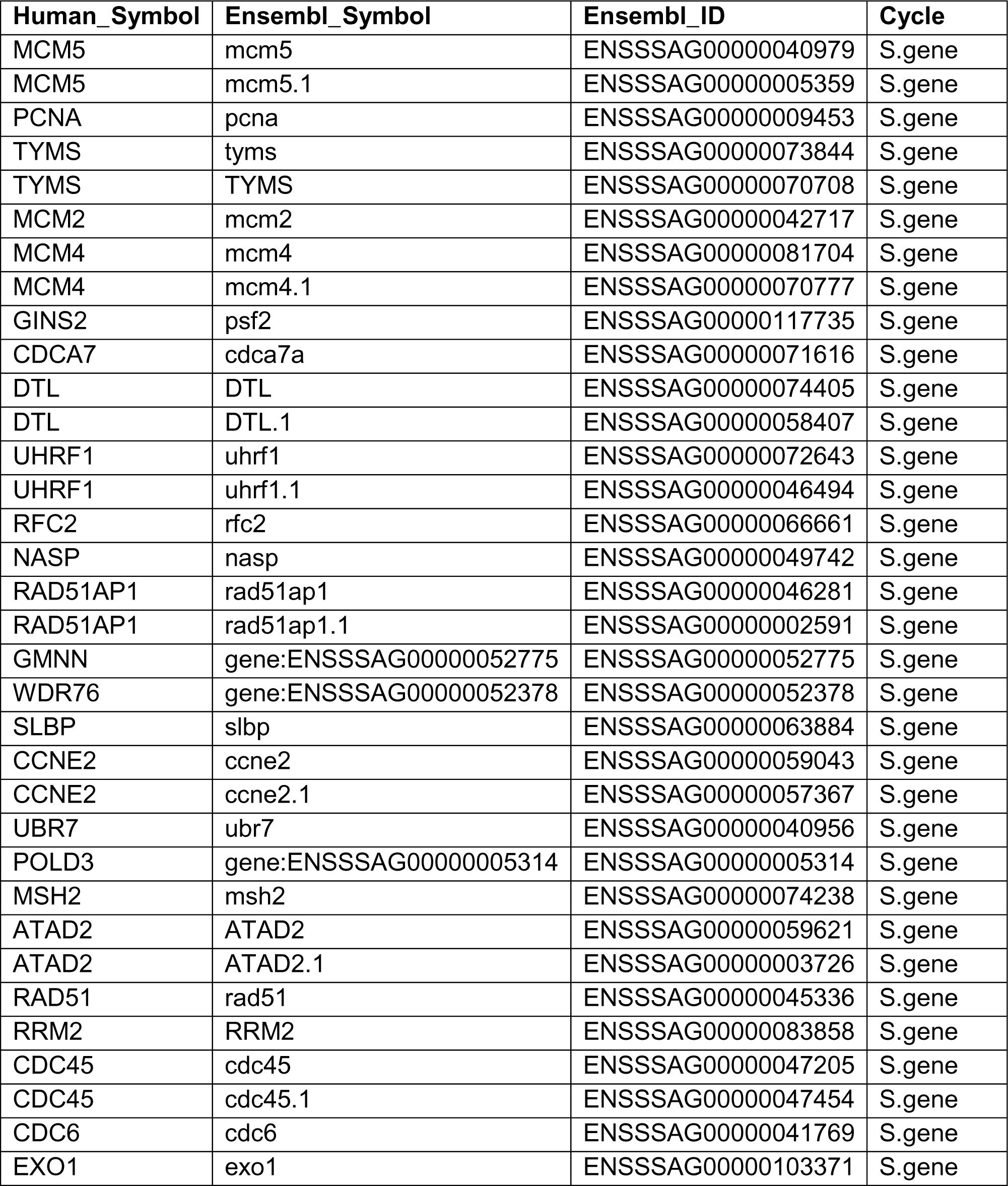
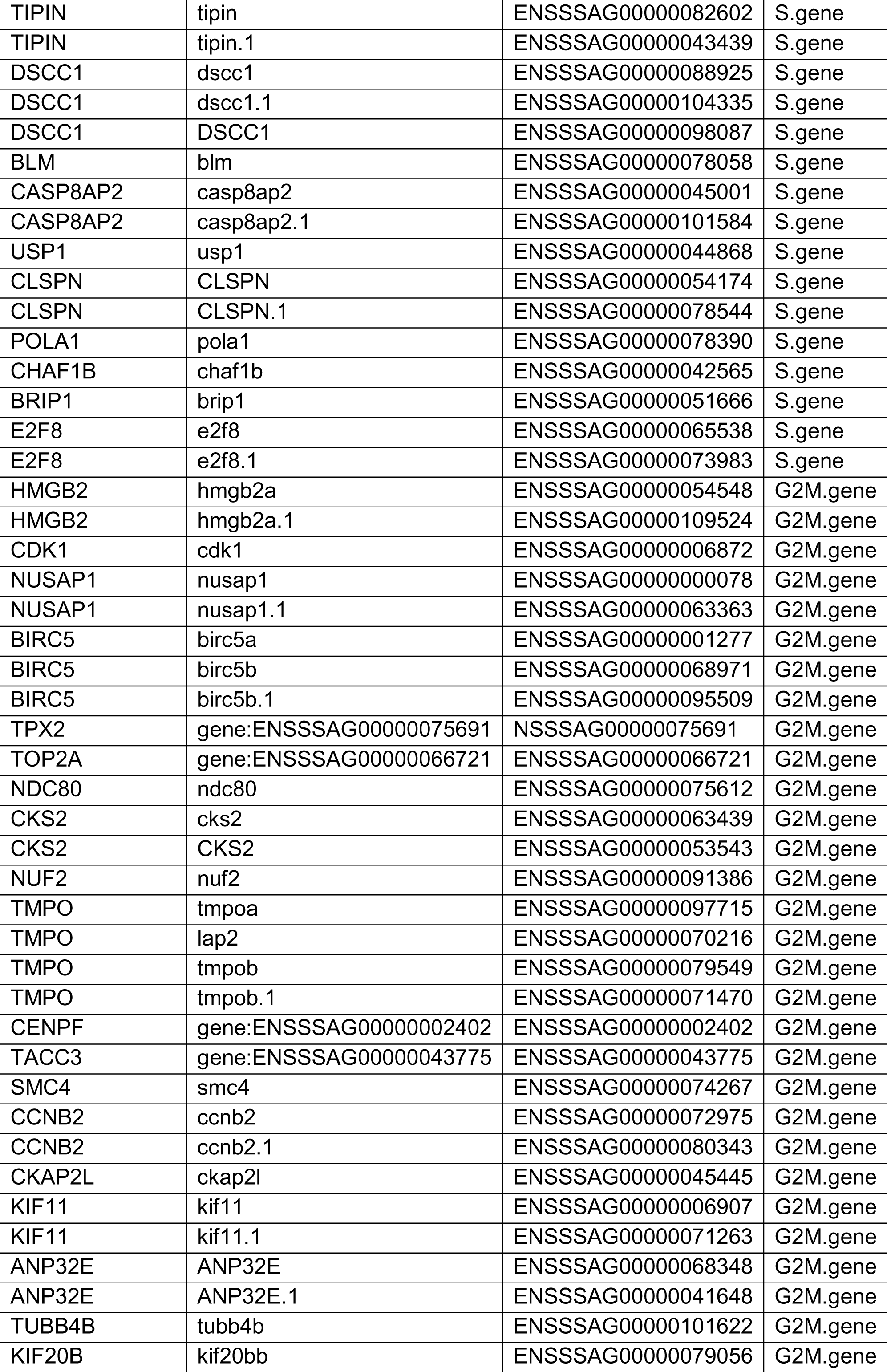
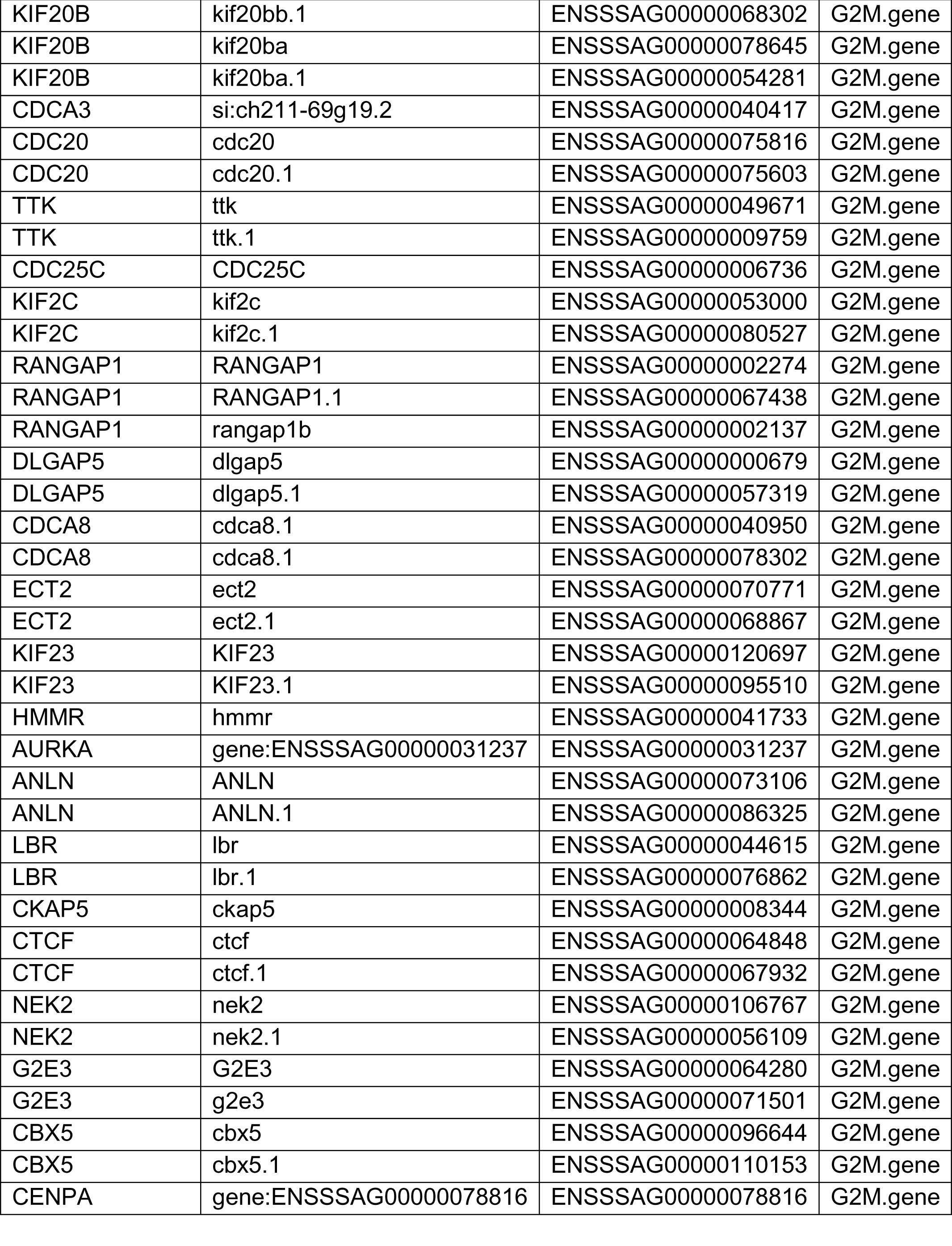
Sequencing statistics for the six different single-cell libraries and four spatial transcriptomic libraries. (In Excell file)

**Supplementary Table 2.** Genes used for cell cycle scoring for Atlantic salmon.

| Sample ID | Cell Count |  |  |  |
| --- | --- | --- | --- | --- |
|  | 1) Nuclei with < 10% of UMIs as mtDNA features | 2) Nuclei passing upper and lower UMI/feature thresholds | 3) Nuclei after removing doublets | 4) Nuclei after removing poor-quality clusters |
| Atlantic_Canada_dorsal_fin | 58906 | 9080 | 8780 | 6420 |
| Atlantic_Canada_pelvic_fin | 55206 | 7896 | 7647 | 4343 |
| Atlantic_Canada_skin | 36325 | 13648 | 13217 | 6874 |
| Atlantic_Scotland_pectoral_fin | 54586 | 9055 | 8760 | 6411 |
| Atlantic_Scotland_pelvic_fin | 61876 | 3057 | 2953 | 2102 |
| Atlantic_Scotland_skin | 52270 | 2223 | 2143 | 1839 |

**Supplementary Table 3.**
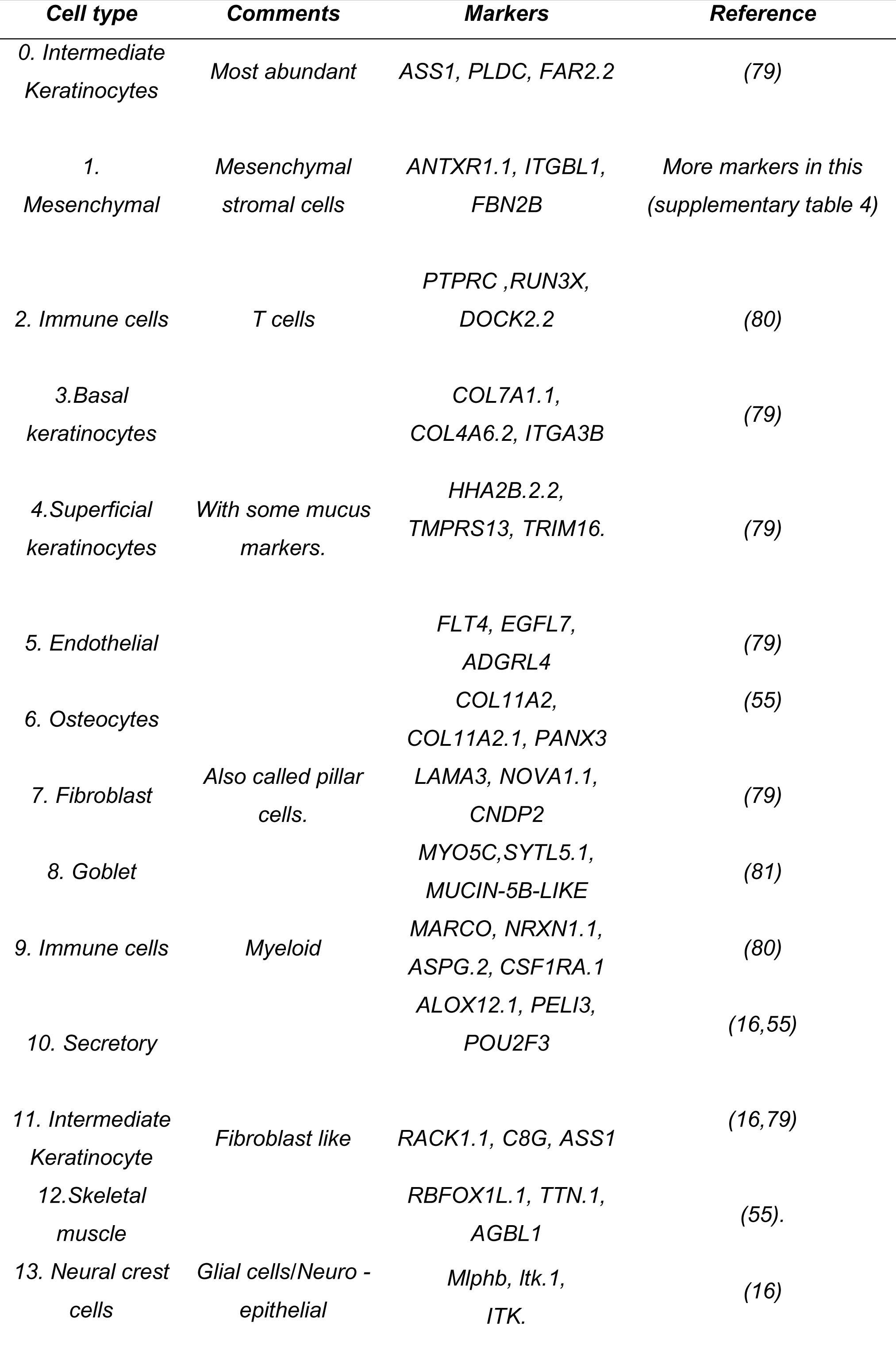
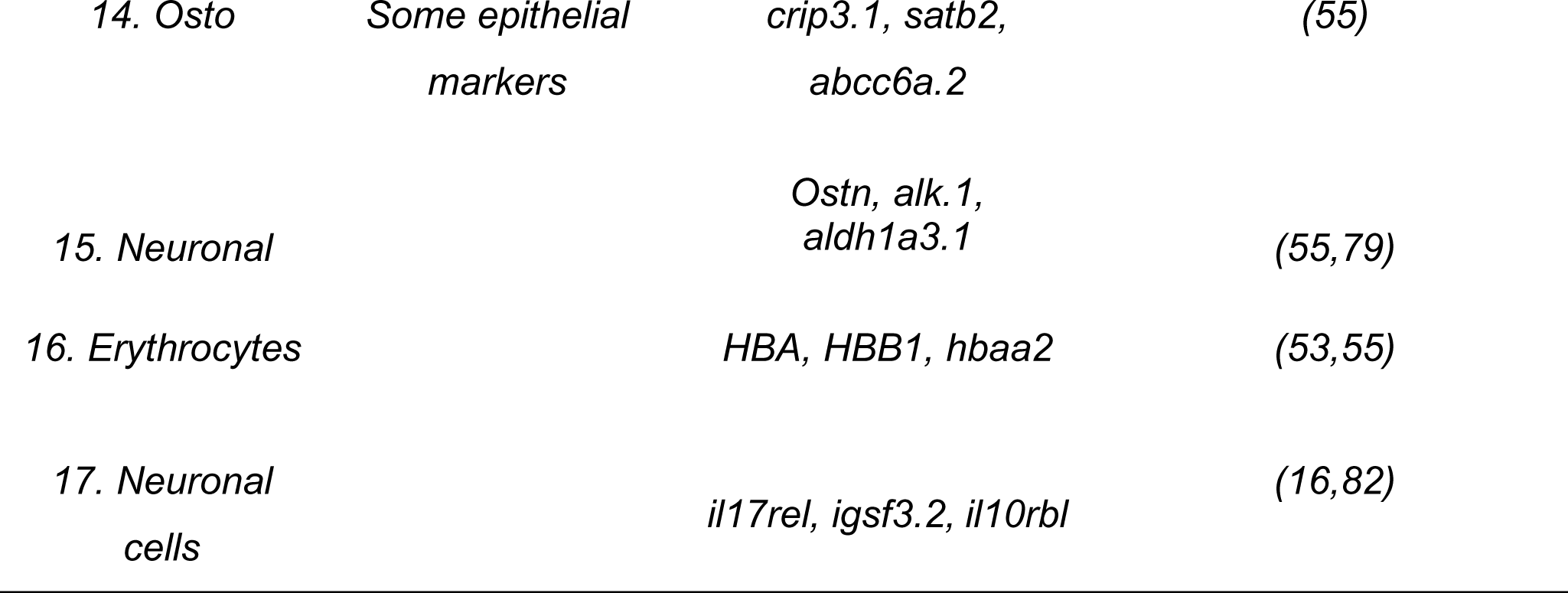
Number of nuclei detected per sample after several filtering stages: 1) removing nuclei where mtDNA features accounted for 10% or more of all UMIs, 2) removing nuclei that exceeded UMI/feature upper and lower bounds, 3) removing doublet cells identified with DoubletFinder, and 4) removing cells in poor-quality clusters.

**Supplementary Table 9** Stromal cell markers for Seurat-generated skin samples. (in Excell file)

**Supplementary Table 10** Stromal cell markers for Seurat clustering. (in Excell file)

## REFERENCES

1. FAO. The State of World Fisheries and Aquaculture 2020: Sustainability in action [Internet]. Rome, Italy: FAO; 2020 [cited 2023 Jun 12]. 244 p. (The State of World Fisheries and Aquaculture (SOFIA)). Available from: https://www.fao.org/documents/card/en/c/ca9229en

2. Norwegian Fish Health Report 2022 [Internet]. [cited 2023 Jun 12]. Available from: https://www.vetinst.no/rapporter-og-publikasjoner/rapporter/2023/norwegian-fish-health-report-2022

3. Melo FR, Bressan RB, Forner S, Martini AC, Rode M, Delben PB, et al. Transplantation of Human Skin-Derived Mesenchymal Stromal Cells Improves Locomotor Recovery After Spinal Cord Injury in Rats. Cell Mol Neurobiol. 2017 Jul;37(5):941–7.

4. Scadden DT. The stem-cell niche as an entity of action. Nature. 2006 Jun 29;441(7097):1075–9.

5. Tumbar T, Guasch G, Greco V, Blanpain C, Lowry WE, Rendl M, et al. Defining the Epithelial Stem Cell Niche in Skin. Science. 2004 Jan 16;303(5656):359–63.

6. Deb A. How Stem Cells Turn into Bone and Fat. N Engl J Med. 2019 Jun 6;380(23):2268–70.

7. Horwitz EM, Andreef M, Frassoni F. Mesenchymal Stromal Cells. Curr Opin Hematol. 2006 Nov;13(6):419–25.

8. Kuroda J, Itabashi T, Iwane AH, Aramaki T, Kondo S. The Physical Role of Mesenchymal Cells Driven by the Actin Cytoskeleton Is Essential for the Orientation of Collagen Fibrils in Zebrafish Fins. Front Cell Dev Biol. 2020 Oct 14;8:580520.

9. Ytteborg E, Todorcevic M, Krasnov A, Takle H, Kristiansen IØ, Ruyter B. Precursor cells from Atlantic salmon (Salmo salar) visceral fat holds the plasticity to differentiate into the osteogenic lineage. Biol Open. 2015 Jul 15;4(7):783–91.

10. Rakers S, Gebert M, Uppalapati S, Meyer W, Maderson P, Sell AF, et al. ‘Fish matters’: the relevance of fish skin biology to investigative dermatology. Exp Dermatol. 2010;19(4):313–24.

11. Slyper M, Porter CBM, Ashenberg O, Waldman J, Drokhlyansky E, Wakiro I, et al. A single-cell and single-nucleus RNA-Seq toolbox for fresh and frozen human tumors. Nat Med. 2020 May 1;26(5):792–802.

12. Andresen AMS, Taylor RS, Grimholt U, Daniels RR, Sun J, Dobie R, et al. Mapping the cellular landscape of Atlantic salmon head kidney by single cell and single nucleus transcriptomics. Fish Shellfish Immunol. 2024 Mar 1;146:109357.

13. Ruiz Daniels R, Taylor RS, Robledo D, Macqueen DJ. Single cell genomics as a transformative approach for aquaculture research and innovation. Rev Aquac. 2023;15(4):1618–37.

14. Marx V. Method of the Year: spatially resolved transcriptomics. Nat Methods. 2021 Jan 6;18(1):9–14.

15. Chen CJ, Kajita H, Takaya K, Aramaki-Hattori N, Sakai S, Asou T, et al. Single-Cell RNA-seq Analysis Reveals Cellular Functional Heterogeneity in Dermis Between Fibrotic and Regenerative Wound Healing Fates. Front Immunol [Internet]. 2022 [cited 2023 Jul 26];13. Available from: 10.3389/fimmu.2022.875407

16. Salisbury SJ, Ruiz Daniels R, Monaghan SJ, Bron JE, Villamayor PR, Gervais O, et al. Keratinocytes Drive the Epithelial Hyperplasia Key to Sea Lice Resistance in Coho Salmon [Internet]. bioRxiv; 2023 [cited 2023 Nov 20]. p. 2023.10.15.562030. Available from: 10.1101/2023.10.15.562030v1

17. Sveen LR, Robinson N, Krasnov A, Ruiz Daniels R, Vaadal M, Karlsen C, et al. Transcriptomic landscape of Atlantic salmon (Salmo salar L.) skin. G3 GenesGenomesGenetics. 2023 Sep 19;jkad215.

18. Togarrati PP, Sasaki RT, Abdel-Mohsen M, Dinglasan N, Deng X, Desai S, et al. Identification and characterization of a rich population of CD34+ mesenchymal stem/stromal cells in human parotid, sublingual and submandibular glands. Sci Rep. 2017 Jun 14;7(1):3484.

19. Cui LL, Nitzsche F, Pryazhnikov E, Tibeykina M, Tolppanen L, Rytkönen J, et al. Integrin α4 Overexpression on Rat Mesenchymal Stem Cells Enhances Transmigration and Reduces Cerebral Embolism After Intracarotid Injection. Stroke. 2017 Oct;48(10):2895– 900.

20. Hamidouche Z, Fromigué O, Ringe J, Häupl T, Vaudin P, Pagès JC, et al. Priming integrin α5 promotes human mesenchymal stromal cell osteoblast differentiation and osteogenesis. Proc Natl Acad Sci U S A. 2009 Nov 3;106(44):18587–91.

21. Chari T, Pachter L. The specious art of single-cell genomics. PLOS Comput Biol. 2023 ago;19(8):e1011288.

22. Moon KR, van Dijk D, Wang Z, Gigante S, Burkhardt DB, Chen WS, et al. Visualizing structure and transitions in high-dimensional biological data. Nat Biotechnol. 2019 Dec;37(12):1482–92.

23. Zhou Y, Liu C, He J, Dong L, Zhu H, Zhang B, et al. KLF2+ stemness maintains human mesenchymal stem cells in bone regeneration. Stem Cells Dayt Ohio. 2020 Mar;38(3):395–409.

24. Liu J, He S, Ma B, Li X, Wang Y, Xiong J. TMT-based quantitative proteomic analysis revealed that FBLN2 and NPR3 are involved in the early osteogenic differentiation of mesenchymal stem cells (MSCs). Aging. 2023 Aug 4;15(15):7637–54.

25. Zhang L, Xie H, Li S. LncRNA LOXL1-AS1 controls osteogenic and adipocytic differentiation of bone marrow mesenchymal stem cells in postmenopausal osteoporosis through regulating the miR-196a-5p/Hmga2 axis. J Bone Miner Metab. 2020 Nov;38(6):794–805.

26. Ambele MA, Dhanraj P, Giles R, Pepper MS. Adipogenesis: A Complex Interplay of Multiple Molecular Determinants and Pathways. Int J Mol Sci. 2020 Jun 16;21(12):4283.

27. Glunk V, Laber S, Sinnott-Armstrong N, Sobreira DR, Strobel SM, Batista TM, et al. A non-coding variant linked to metabolic obesity with normal weight affects actin remodelling in subcutaneous adipocytes. Nat Metab. 2023 May;5(5):861–79.

28. Kubo H, Shimizu M, Taya Y, Kawamoto T, Michida M, Kaneko E, et al. Identification of mesenchymal stem cell (MSC)-transcription factors by microarray and knockdown analyses, and signature molecule-marked MSC in bone marrow by immunohistochemistry. Genes Cells Devoted Mol Cell Mech. 2009 Mar;14(3):407–24.

29. Wei K, Xu Y, Tse H, Manolson MF, Gong SG. Mouse FLRT2 interacts with the extracellular and intracellular regions of FGFR2. J Dent Res. 2011 Oct;90(10):1234–9.

30. Martin TJ. PTH1R Actions on Bone Using the cAMP/Protein Kinase A Pathway. Front Endocrinol [Internet]. 2022 [cited 2024 Feb 2];12. Available from: 10.3389/fendo.2021.833221

31. Bae JH, Park D. Effect of dietary calcium on the gender-specific association between polymorphisms in the PTPRD locus and osteoporosis. Clin Nutr Edinb Scotl. 2022 Mar;41(3):680–6.

32. Qin X, Jiang Q, Matsuo Y, Kawane T, Komori H, Moriishi T, et al. Cbfb regulates bone development by stabilizing Runx family proteins. J Bone Miner Res Off J Am Soc Bone Miner Res. 2015 Apr;30(4):706–14.

33. Wang Y, Feng Q, Ji C, Liu X, Li L, Luo J. RUNX3 plays an important role in mediating the BMP9-induced osteogenic differentiation of mesenchymal stem cells. Int J Mol Med. 2017 Dec;40(6):1991–9.

34. Mucientes A, Herranz E, Moro E, González-Corchón A, Peña-Soria MJ, Abasolo L, et al. Influence of Mesenchymal Stem Cell Sources on Their Regenerative Capacities on Different Surfaces. Cells. 2021 Feb 23;10(2):481.

35. Kim JM, Lin C, Stavre Z, Greenblatt MB, Shim JH. Osteoblast-Osteoclast Communication and Bone Homeostasis. Cells. 2020 Sep 10;9(9):2073.

36. Bagchi DP, Li Z, Corsa CA, Hardij J, Mori H, Learman BS, et al. Wntless regulates lipogenic gene expression in adipocytes and protects against diet-induced metabolic dysfunction. Mol Metab. 2020 Sep;39:100992.

37. Kislev N, Mor-Yossef Moldovan L, Barak R, Egozi M, Benayahu D. MYH10 Governs Adipocyte Function and Adipogenesis through Its Interaction with GLUT4. Int J Mol Sci. 2022 Feb 21;23(4):2367.

38. Thomas T, Gori F, Khosla S, Jensen MD, Burguera B, Riggs BL. Leptin acts on human marrow stromal cells to enhance differentiation to osteoblasts and to inhibit differentiation to adipocytes. Endocrinology. 1999 Apr;140(4):1630–8.

39. Wang Y, Chen D, Zhang Y, Wang P, Zheng C, Zhang S, et al. Novel Adipokine, FAM19A5, Inhibits Neointima Formation After Injury Through Sphingosine-1-Phosphate Receptor 2. Circulation. 2018 Jul 3;138(1):48–63.

40. Menssen A, Häupl T, Sittinger M, Delorme B, Charbord P, Ringe J. Differential gene expression profiling of human bone marrow-derived mesenchymal stem cells during adipogenic development. BMC Genomics. 2011 Sep 24;12(1):461.

41. Zaman G, Staines KA, Farquharson C, Newton PT, Dudhia J, Chenu C, et al. Expression of Sulf1 and Sulf2 in cartilage, bone and endochondral fracture healing. Histochem Cell Biol. 2016 Jan;145(1):67–79.

42. Nassari S, Duprez D, Fournier-Thibault C. Non-myogenic Contribution to Muscle Development and Homeostasis: The Role of Connective Tissues. Front Cell Dev Biol. 2017;5:22.

43. von Hofsten J, Elworthy S, Gilchrist MJ, Smith JC, Wardle FC, Ingham PW. Prdm1- and Sox6-mediated transcriptional repression specifies muscle fibre type in the zebrafish embryo. EMBO Rep. 2008 Jul;9(7):683–9.

44. Zhao S, Zhao Y, Niu P, Wang N, Tang Z, Zan L, et al. Molecular Characterization of Porcine MMP19 and MMP23B Genes and Its Association with Immune Traits. Int J Biol Sci. 2011 Sep 14;7(8):1101–13.

45. Moon H, Zhu J, Donahue LR, Choi E, White AC. Krt5+/Krt15+ foregut basal progenitors give rise to cyclooxygenase-2-dependent tumours in response to gastric acid stress. Nat Commun. 2019 May 20;10(1):2225.

46. Corcoran JP, Ferretti P. Keratin 8 and 18 expression in mesenchymal progenitor cells of regenerating limbs is associated with cell proliferation and differentiation. Dev Dyn Off Publ Am Assoc Anat. 1997 Dec;210(4):355–70.

47. Bou M, Montfort J, Le Cam A, Rallière C, Lebret V, Gabillard JC, et al. Gene expression profile during proliferation and differentiation of rainbow trout adipocyte precursor cells. BMC Genomics. 2017 May 4;18(1):347.

48. Jimenez MA, Åkerblad P, Sigvardsson M, Rosen ED. Critical Role for Ebf1 and Ebf2 in the Adipogenic Transcriptional Cascade. Mol Cell Biol. 2007 Jan;27(2):743–57.

49. Bobowski-Gerard M, Boulet C, Zummo FP, Dubois-Chevalier J, Gheeraert C, Bou Saleh M, et al. Functional genomics uncovers the transcription factor BNC2 as required for myofibroblastic activation in fibrosis. Nat Commun. 2022 Sep 10;13(1):5324.

50. Tsai HL, Chiu WT, Fang CL, Hwang SM, Renshaw PF, Lai WFT. Different Forms of Tenascin-C with Tenascin-R Regulate Neural Differentiation in Bone Marrow-Derived Human Mesenchymal Stem Cells. Tissue Eng Part A. 2014 Jul 1;20(13–14):1908–21.

51. Alcaraz LB, Exposito JY, Chuvin N, Pommier RM, Cluzel C, Martel S, et al. Tenascin-X promotes epithelial-to-mesenchymal transition by activating latent TGF-β. J Cell Biol. 2014 May 12;205(3):409–28.

52. Wu H, Ma S, Xiang M, Tong S. HTRA1 promotes transdifferentiation of normal fibroblasts to cancer-associated fibroblasts through activation of the NF-κB/bFGF signaling pathway in gastric cancer. Biochem Biophys Res Commun. 2019 Jun 30;514(3):933–9.

53. ZFIN The Zebrafish Information Network [Internet]. [cited 2023 Nov 28]. Available from: https://zfin.org/

54. Parente R, Sobacchi C, Bottazzi B, Mantovani A, Grčevic D, Inforzato A. The Long Pentraxin PTX3 in Bone Homeostasis and Pathology. Front Immunol. 2019 Nov 8;10:2628.

55. Farnsworth DR, Saunders LM, Miller AC. A single-cell transcriptome atlas for zebrafish development. Dev Biol. 2020 Mar 15;459(2):100–8.

56. Stosik M, Tokarz-Deptuła B, Deptuła J, Deptuła W. Immune Functions of Erythrocytes in Osteichthyes. Front Immunol. 2020 Sep 15;11:1914.

57. Rasmussen JP, Vo NT, Sagasti A. Fish Scales Dictate the Pattern of Adult Skin Innervation and Vascularization. Dev Cell. 2018 Aug;46(3):344–359.e4.

58. Sehring I, Mohammadi HF, Haffner-Luntzer M, Ignatius A, Huber-Lang M, Weidinger G. Zebrafish fin regeneration involves generic and regeneration-specific osteoblast injury responses. eLife. 11:e77614.

59. Ugurlu B, Karaoz E. Comparison of similar cells: Mesenchymal stromal cells and fibroblasts. Acta Histochem. 2020 Dec;122(8):151634.

60. Soundararajan M, Kannan S. Fibroblasts and mesenchymal stem cells: Two sides of the same coin? J Cell Physiol. 2018;233(12):9099–109.

61. Augello A, De Bari C. The regulation of differentiation in mesenchymal stem cells. Hum Gene Ther. 2010 Oct;21(10):1226–38.

62. Richardson R, Slanchev K, Kraus C, Knyphausen P, Eming S, Hammerschmidt M. Adult Zebrafish as a Model System for Cutaneous Wound-Healing Research. J Invest Dermatol. 2013 Jun 1;133(6):1655–65.

63. Richardson R, Metzger M, Knyphausen P, Ramezani T, Slanchev K, Kraus C, et al. Re- epithelialization of cutaneous wounds in adult zebrafish combines mechanisms of wound closure in embryonic and adult mammals. Development. 2016 Jun 15;143(12):2077–88.

64. Maxson S, Lopez EA, Yoo D, Danilkovitch-Miagkova A, LeRoux MA. Concise Review: Role of Mesenchymal Stem Cells in Wound Repair. Stem Cells Transl Med. 2012 Feb;1(2):142–9.

65. Novoseletskaya E, Grigorieva O, Nimiritsky P, Basalova N, Eremichev R, Milovskaya I, et al. Mesenchymal Stromal Cell-Produced Components of Extracellular Matrix Potentiate Multipotent Stem Cell Response to Differentiation Stimuli. Front Cell Dev Biol [Internet]. 2020 [cited 2024 Feb 4];8. Available from: 10.3389/fcell.2020.555378

66. Song N, Scholtemeijer M, Shah K. Mesenchymal Stem Cell Immunomodulation: Mechanisms and Therapeutic potential. Trends Pharmacol Sci. 2020 Sep;41(9):653–64.

67. Sveen LR, Timmerhaus G, Krasnov A, Takle H, Handeland S, Ytteborg E. Wound healing in post-smolt Atlantic salmon (Salmo salar L.). Sci Rep. 2019 Mar 5;9(1):3565.

68. Tencerova M, Frost M, Figeac F, Nielsen TK, Ali D, Lauterlein JJL, et al. Obesity- Associated Hypermetabolism and Accelerated Senescence of Bone Marrow Stromal Stem Cells Suggest a Potential Mechanism for Bone Fragility. Cell Rep. 2019 May;27(7):2050–2062.e6.

69. Rauch A, Haakonsson AK, Madsen JGS, Larsen M, Forss I, Madsen MR, et al. Osteogenesis depends on commissioning of a network of stem cell transcription factors that act as repressors of adipogenesis. Nat Genet. 2019 Apr;51(4):716–27.

70. Drokhlyansky E, Smillie CS, Van Wittenberghe N, Ericsson M, Griffin GK, Eraslan G, et al. The Human and Mouse Enteric Nervous System at Single-Cell Resolution. Cell. 2020 Sep 17;182(6):1606–1622.e23.

71. Daniels RR, Taylor RS, Dobie R, Salisbury S, Furniss JJ, Clark E, et al. A versatile nuclei extraction protocol for single nucleus sequencing in non-model species– Optimization in various Atlantic salmon tissues. PLOS ONE. 2023 Sep 7;18(9):e0285020.

72. Dobin A, Davis CA, Schlesinger F, Drenkow J, Zaleski C, Jha S, et al. STAR: Ultrafast universal RNA-seq aligner. Bioinformatics. 2013 Jan;29(1):15–21.

73. Pertea G, Pertea M. GFF Utilities: GffRead and GffCompare. F1000Research. 2020;9:ISCB Comm J-304.

74. Hao Y, Hao S, Andersen-Nissen E, Mauck WM, Zheng S, Butler A, et al. Integrated analysis of multimodal single-cell data. Cell. 2021 Jun 24;184(13):3573–3587.e29.

75. Ahlmann-Eltze C, Huber W. glmGamPoi: fitting Gamma-Poisson generalized linear models on single cell count data. Bioinformatics. 2021 Apr 5;36(24):5701–2.

76. Mcginnis CS, Murrow LM, Correspondence ZJG. DoubletFinder: Doublet Detection in Single-Cell RNA Sequencing Data Using Artificial Nearest Neighbors. Cell Syst. 2019;8:329–337.e4.

77. Al-Obaide M, Ishmakej A, Brown C, Mazzella M, Agosta P, Perez-Cruet M, et al. The potential role of integrin alpha 6 in human mesenchymal stem cells. Front Genet. 2022 Sep 16;13:968228.

78. Nieto-Nicolau N, de la Torre RM, Fariñas O, Savio A, Vilarrodona A, Casaroli-Marano RP. Extrinsic modulation of integrin α6 and progenitor cell behavior in mesenchymal stem cells. Stem Cell Res. 2020 Aug 1;47:101899.

79. The Human Protein Atlas [Internet]. [cited 2024 Feb 19]. Available from: https://www.proteinatlas.org/

80. Taylor RS, Ruiz Daniels R, Dobie R, Naseer S, Clark TC, Henderson NC, et al. Single cell transcriptomics of Atlantic salmon (Salmo salar L.) liver reveals cellular heterogeneity and immunological responses to challenge by Aeromonas salmonicida. Front Immunol [Internet]. 2022 [cited 2022 Sep 18];13. Available from: 10.3389/fimmu.2022.984799

81. West AC, Mizoro Y, Wood SH, Ince LM, Iversen M, Jørgensen EH, et al. Immunologic Profiling of the Atlantic Salmon Gill by Single Nuclei Transcriptomics. Front Immunol. 2021;12:1.

82. Tambalo M, Mitter R, Wilkinson DG. A single cell transcriptome atlas of the developing zebrafish hindbrain. Development. 2020 Mar 16;147(6):dev184143.

